# “Universal Hyb-Seq kits capture considerable intraspecific variation: Less is more in herbarium-inclusive molecular ecology”

**DOI:** 10.1101/2025.10.14.682325

**Authors:** Yannick Woudstra, Anne-Sophie Quatela, Niels CAM Wagemaker, Slavica Ivanovic, Tanja Slotte, Patrick Meirmans, Barbara Gravendeel, Koen JF Verhoeven

## Abstract

Target capture sequencing has enhanced the study of plant evolution and molecular ecology, particularly through the access to degraded DNA from herbarium specimens. Universal “off-the-shelf” kits, such as Angiosperms-353, are cheap and readily available but are considered to expose insufficient variation below the species level, because they are designed to target highly conserved regions. However, this remains to be tested in a direct comparison with customised approaches below the species level. In this study, near-identical genotypes from both herbarium and fresh material of the common dandelion (apomictic lineages in *Taraxacum officinale* F.H.Wigg.) are characterised with customised and universal approaches of target capture sequencing. An RNA-bait panel was designed to capture (i) highly variable loci normally obtained with a Genotyping-by-Sequencing (GBS) approach customised for dandelions; (ii) custom selected genes with potential for environmental adaptation, likely to harbour intraspecific genetic variation; (iii) conserved exons from universal kits (Angiosperms-353; Compositae-COS). Although exons from universal kits yield considerably less intraspecific genetic variation than both customised approaches, they still provided sufficient genetic variation to discriminate between near-identical genotypes of the same apomictic lineage. Given that universal kits save time, money, and the need for genomic reference data, this approach is recommended to increase the number of samples under budgetary constraints while still capturing considerable levels of intraspecific genetic variation.

## Introduction

Herbarium genomics, the ability to access sequence information from historical voucher specimens, has revolutionised studies of plant evolution and molecular ecology (Bakker et al. 2020). Herbarium collections provide access to expertly identified material of virtually all plant species ever described, making them particularly useful for studies of molecular systematics (Dodsworth et al. 2019). Below the species level, they represent unique records of historical populations, giving insights into historical floras (Slimp et al. 2021), extinct populations (Rosche et al. 2022) and evolutionary responses to climate change (Lang et al. 2024). The (generally) degraded nature of DNA from historical samples makes it, however, challenging to obtain this historical genomic information in a cost-effective way. To improve the efficiency in upscaling herbarium-inclusive molecular ecology, a critical evaluation of existing tools is needed.

Optimising the cost-efficiency of obtaining informative genetic markers has been a key challenge in population genomics for the last decades, in search of increasing the number of samples (Hale et al. 2020). Although neutral markers such as AFLPs and microsatellites are cost-effective in elucidating demographic patterns, they generally do not reflect differential selection pressures (Narum et al. 2013). Whole-genome sequencing of hundreds of samples may be feasible for model plants with small genomes (e.g., relatives of *Arabidopsis thaliana*) but is still unrealistic with larger genomes, particularly in the absence of (closely related) reference genome assemblies. Restriction-based reduced representation sequencing, such as RADseq or Genotyping-by-Sequencing (GBS), is therefore a popular alternative as it captures variable genome-wide markers with limited sequencing efforts, allowing multiplexed sequencing of hundreds of individuals (Elshire et al. 2011). By digesting genomic DNA samples with sequence-specific restriction enzymes followed by size selection on small fragments, the sequencing library is reduced to a subsample of the genome, generating thousands of meaningful SNPs, even for very large genomes (e.g. conifers, Pan et al. 2015) and in non-model organisms (e.g. *Reaumuria trigyna*, Dang et al. 2024). The technique is, however, less suitable for historical (herbarium) samples as it relies on the digestion of large intact genomic fragments, which are rare in the generally degraded DNA obtained from historical samples (Staats et al. 2011).

Target capture sequencing has become the go-to sequencing tool when working with historical (degraded) DNA samples, such as herbarium material (Dodsworth et al. 2019). Through the use of short (80-120 bases) oligonucleotide baits with sequences complementary to target genes, target DNA fragments are captured to effectively reduce the complexity of sequencing libraries to include only loci of interest (Gnirke et al. 2009). The use of sequencing power in obtaining sequence data for (low-copy) nuclear genes is thereby optimised. This circumvents the limitations of PCR-based targeted sequencing, such as amplicon sequencing, or restriction-based reduced representation techniques, such as GBS, which rely on long uninterrupted DNA fragments (Woudstra et al. 2022). Although target capture applications of restriction-based approaches do exist (e.g., Barreiro et al. 2017; Lang et al. 2020), the set-up is time-consuming, expensive and has rarely been tested in groups with large phylogenetic distance to model organisms (e.g., *Arabidopsis thaliana*). Target capture studies therefore usually focus on (low-copy) nuclear genes, for which reference sequences can readily be obtained through transcriptome resources, even from related clades (Woudstra et al. 2022).

A popular and upscaleable approach in herbarium genomics is the use of universal target capture kits. These kits can be used at broad taxonomic ranges, such as all flowering plants (Angiosperms-353; Johnson et al. 2019), are reasonably priced, and readily available. The downside is that they target highly conserved exonic regions, limiting their application below the species level. Higher taxonomic resolution is generally obtained with a customised target capture approach, targeting exons from more variable genes (e.g. *Aloe*, Woudstra et al. 2021; Orchidiinae, Veltman et al. 2024). Customisation also allows the targeting of more variable intronic regions (Quatela et al. 2025), as well as genes related to local adaptation (e.g. shade adaptation, Michel et al. 2022). Customised kits have shown great potential below the species level, with impactful applications in tracing international plant trade (*Anacyclus pyrethrum*, Manzanilla et al. 2022; *Aloe vera*, Woudstra et al. 2025). The required investment of time and resources (i.e. generating genomic reference data), however, makes a customised approach less suitable for upscaling to hundreds of samples.

Amid growing evidence that universal kits can capture sufficient intraspecific variation in some clades (e.g., 24 Texan plant species, Slimp et al. 2021; *Solidago ulmifolia*, Beck et al. 2021; *Dactylorhiza cantabrica*, Otero *et al*. 2024; several tree species, Ousmael and Hansen 2025; *Androsace cantabrica*, Liang et al. 2025), the question is raised whether investment in a customised kit is necessary for intraspecific herbarium genomics. Direct comparisons between custom and universal kits in the same species are scarce (Yardeni et al. 2022) and lack a definitive answer on differences in obtained intraspecific resolution.

This study presents a direct comparison of discriminative power between loci from customised and universal target capture kits for use with both fresh and herbarium samples (Fig. 1). For this we use a uniquely suitable system for such a comparison: apomictic clonal lineages of the common dandelion (*Taraxacum officinale*). Because of the asexual reproduction of dandelions, the genotypes of clonal lineages from herbarium specimens and fresh specimens are expected to be nearly identical, which offers a great opportunity to verify the accuracy of different methods for genotyping herbarium samples. By sequencing verified samples from present and historical collections of these clonal lineages, the limits of herbarium-inclusive molecular ecology are tested and the added value of customised target capture kits is verified.

**Figure 1:**
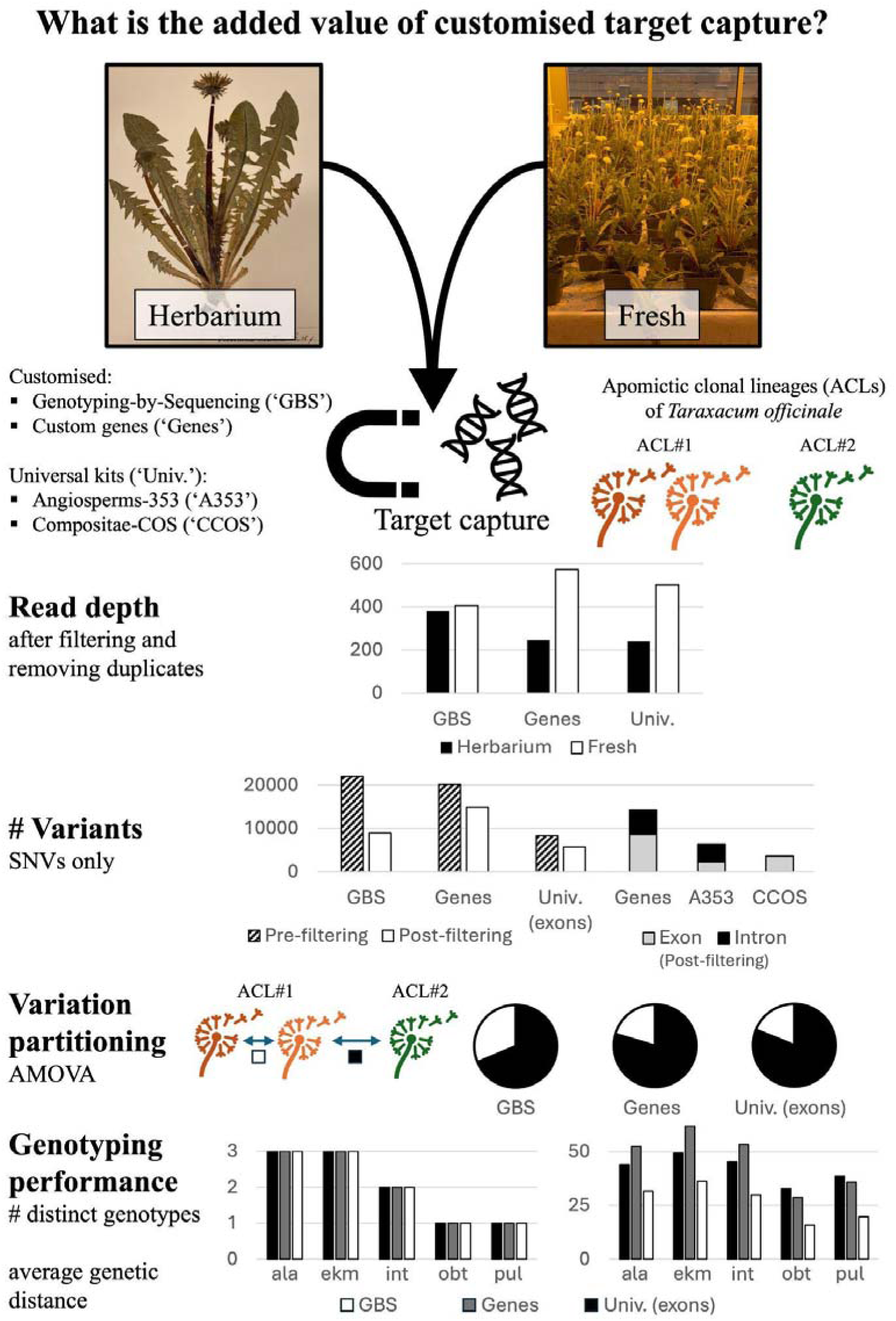
Overview of the study, summarising the main findings. The added value of customised target capture sequencing (manually selecting informative loci) over universal target capture kits is tested in a system of apomictic clonal lineages (ACLs) of dandelions (*Taraxacum officinale*). The genotyping performance for distinguishing differences between ACLs and between individual members of the same ACL (near-identical genotypes) is investigated. Although exons from universal kits yield a considerably lower amount of variation than customised approaches, the variation captured is supported by high read depth and can still be used to effectively characterise all genotypes. Higher resolution is achieved with customised approaches, and (selectively neutral) GBS loci are better at distinguishing near-identical genotypes (i.e. ACLs *obtusifrons* and *pulchrifolium*). The five ACLs investigated in this study are abbreviated to three-letter codes in this figure: ala=*alatum*, ekm=*ekmanii*, int=*interveniens*, obt=*obtusifrons*, pul=*pulchrifolium*. Photo credits: Yannick Woudstra.

## Methods

### 1. Sampling design

Genotyping performance of loci from universal and customised target capture kits was tested in an apomictic complex of dandelions (*Taraxacum officinale* F.H.Wigg.). *T. officinale* is an apomictic species complex with hundreds of clonal lineages that are often derived from mixed sexual-apomictic populations (Kirschner and Štěpánek 1996). Differences between these clonal lineages thus reflect genetic variation found within these mixed populations at the time of origin of the lineages. Although many *Taraxacum* clonal lineages are geographically widespread, their individual members differ only by postpartum somaclonal mutations that have accumulated since their origin (Richards 1996), thereby representing one of the lowest characterisable intraspecific variation levels.

Genotyping using neutral genetic markers (microsatellites) has shown that morphologically recognisable lineages of dandelions (also termed microspecies) are indeed apomictic lineages that are derived from a single clonal founder (Kirschner et al. 2016). For some of these lineages, restriction-based Genotyping-by-Sequencing data is also available (Ibañez et al. 2023), making it possible to develop a customised GBS target capture approach (Lang et al. 2020). Conveniently, verified herbarium material for *Taraxacum* apomictic microspecies is abundant in European botanical collections (i.e. Dutch National Herbarium at Naturalis Biodiversity Center and regional Herbarium Frisicum) thanks to vast collection efforts by European *Taraxacum* aficionados in the 20th century. Altogether, these aspects make *T. officinale* an ideal model system to verify the genotyping limits of custom and universal target capture kits in herbarium genomics.

For comparison in the current study, herbarium and fresh material was sampled from a set of five geographically widespread (Fig. 2) apomictic clonal lineages (ACLs) belonging to *T. officinale.* More specifically, these are the microspecies named *alatum*, *ekmanii*, *interveniens*, *obtusifrons*, and *pulchrifolium*. For each ACL, seeds from 3-4 accessions were selected from the original microsatellite study (Kirschner et al. 2016) for the production of fresh leaf material. These accessions were supplemented with 1-3 historical samples for each ACL, collected between 1954-2000 across central and northern Europe (Fig. 2). Samples came from expertly identified herbarium material from the specialist *Taraxacum* collections of Oosterveld, Van Soest and Hagendijk (Dutch National Herbarium, Naturalis Biodiversity Center). Details on individual accessions can be found in the ‘Accession Information’ file (Online supporting material).

**Figure 2:**
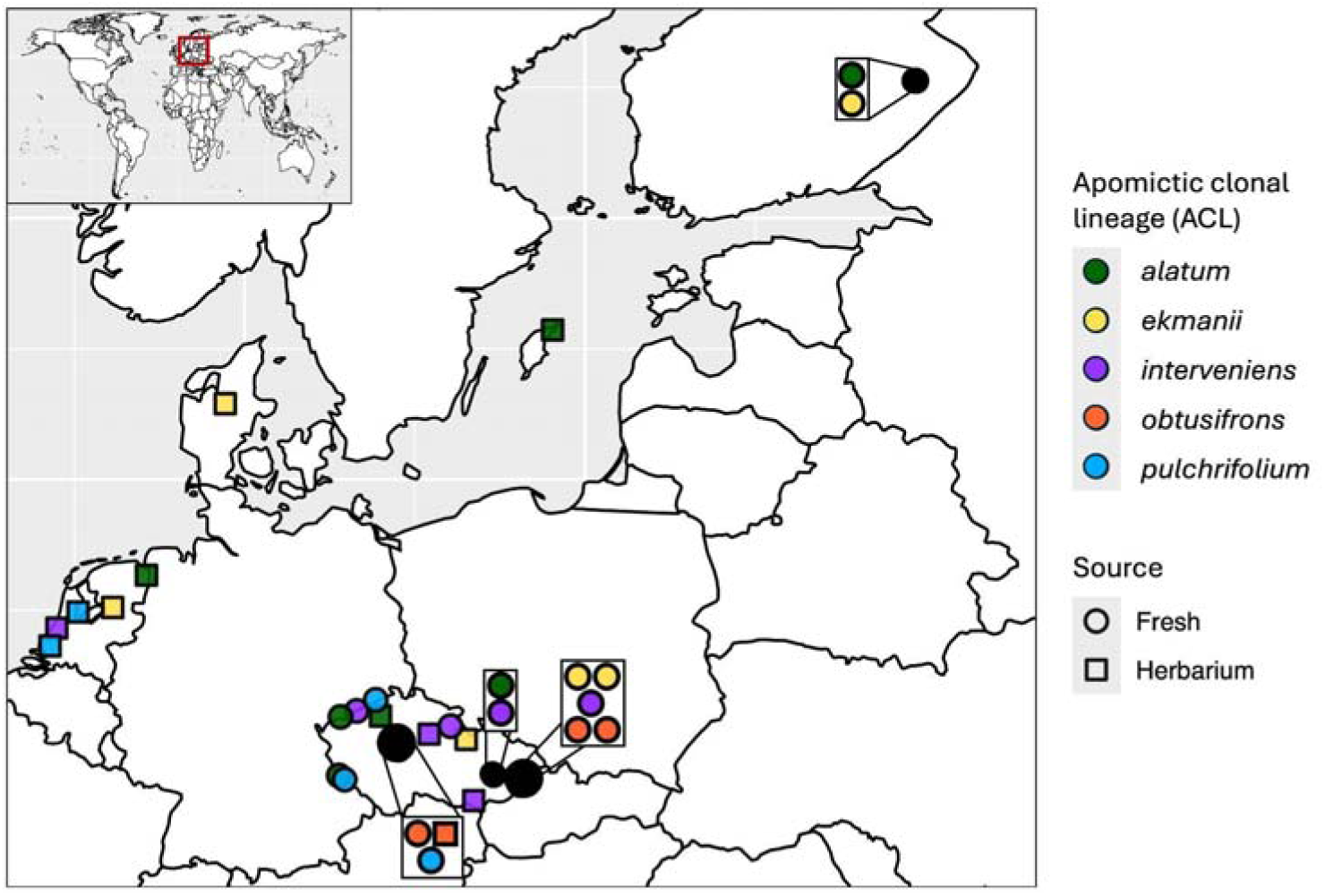
Distribution map of the samples belonging to five widespread apomictic clonal lineages (ACLs) of *Taraxacum officinale* F.H.Wigg., as characterised by Kirschner et al. (2016) using microsatellite data. ACL identity is indicated with different colours, and sample source (herbarium or fresh) is indicated with different symbols. In cases where more than one sample was taken from the same location, sample details are given in expanded boxes. Details on sample provenance, including GPS coordinates, can be found in online supporting material – ‘Accession Information’.

### 2. Target capture development

A target capture sequencing tool was developed to sequence sets of loci with three different levels of expected variation within species. In descending order of expected variation these were: (a) Genotyping-by-Sequencing (GBS) loci; (b) Custom selected nuclear genes with adaptive potential; (c) Nuclear genes captured by two different universal kits [Angiosperms-353 (Johnson et al. 2019) and COS-1061 for Asteraceae (Mandel et al. 2014)]. The protocol for the selection of candidate loci in each category is detailed in dedicated sections a-c below. After selection of nucleotide or (in the case of category b) amino acid sequences, the homologous sequence in *T. officinale* was identified through BLAST searches (Altschul et al. 1990) against the “v11” genome assembly (Xiong et al. 2023). Top hits for each locus or individual exon (categories b and c) were extracted from the BLAST results and transformed into BED files using a python script (Ter Hoeven 2015). For nuclear genes (categories b and c), the individual records within BED files were merged using BEDtools (Quinlan and Hall 2010) to incorporate intronic regions in the target capture design. Dedicated intron-specific BED files were generated from the exon-specific BED files using a custom python script (“make_intron_bed.py”, online supporting material). Loci shorter than 120 bases were discarded to keep target loci at the minimum length of one bait (120 bp). To reduce the risk of capturing high-copy loci, loci occurring on >3 scaffolds with their full (exonic) sequence were discarded. After selecting all suitable loci, an overlap analysis was conducted based on location coordinates in the “v11” genome assembly (BED files) and duplications were removed. In the case of partial overlap, the shorter unique sequence was merged to the larger one.

The target capture tool was developed with the myBaits^®^ protocol (Daicel Arbor Biosciences, Ann Arbor, MI, USA). The selected loci were submitted to Daicel Arbor Biosciences for *in silico* RNA-bait design, where all loci were checked for overlap with known repetitive elements and organellar genome sequences, available in their database (“myBaits^®^ Custom” protocol, Daicel Arbor Biosciences, Ann Arbor, MI, USA). The final set of fully approved loci was translated into a set of 18,346 baits for all three levels of expected variation, targeting 1,458,834 bp of the nuclear genome without tiling (i.e. all nucleotides are targeted by one bait). More details can be found in the online supporting material: full details of the bait design process in the “Target locus selection pipeline” script and “Bioinformatic logbook” files; exon-intron boundaries in the relevant “exon/intron.bed” files; details on loci included in the final bait panel in the ‘Taraxacum target capture design – loci overview’ file.

#### a. GBS loci

The first set of loci was derived from sequencing data from a previous study that used restriction-based Genotyping-by-Sequencing (GBS) to characterise fine-scale genotype differentiation among 20 geographically widespread accessions of the apomictic dandelion lineage *alatum* using a combination of *Csp6I* and *NsiI* restriction enzymes (Ibañez et al., 2023). The raw sequence data (see supplement of Ibañez et al., 2023) was assembled into short (up to 300 bases) GBS loci, using the epiGBS2 pipeline (Gawehns et al., 2022). A total of 2839 loci were included in the target capture design that satisfied each of the following conditions: (1) a minimum read depth of 10, (2) a maximum read depth of 40, (3) recovery in a minimum of 12 samples (out of 20).

#### b. Nuclear genes with adaptive potential

The customised set of full gene loci was developed by conducting an in-depth literature search to identify nuclear genes with evolutionary adaptive potential in dandelions and other angiosperms. Functional genes were chosen based on potential for environmental or ecological adaptation or association with the lineages’ transition from sexual to apomictic reproduction. The 212 loci that were included in the target capture design can be divided into five categories: (I) Apomixis (2 loci); (II) Flowering time regulation (43 genes); (III) Heat resistance (39 genes); (IV) Pollen development (105 genes); and (V) Stomatal regulation (23 genes).

Category I comprises loci controlling two different steps of the apomixis reproductive pathway in *T. officinale*: diplospory (the formation of unreduced megaspores; van Dijk and Bakx-Schotman 2004), and parthenogenesis (the autonomous development of egg cells into embryos without fertilisation; Underwood et al. 2022). As the DIP allele is an unannotated genomic locus, the apomixis loci were not divided into exons and introns.

Flowering time regulation genes (category II), were obtained from the FLOR-ID interactive database on flowering control in *Arabidopsis thaliana* (http://www.flor-id.org; Bouché et al. 2015; accessed in August 2023). Amino acid sequences were obtained from the UniProt (https://www.uniprot.org/) database using accessions listed in FLOR-ID. The explored pathways comprise photoperiod sensitivity, temperature sensitivity, vernalisation, and general control.

For the remaining categories of genes (III to V), UniProt accessions were obtained from literature searches on Google Scholar and specific searches in the AMIGO2 Gene Onthology database (https://amigo.geneontology.org/amigo; accessed in August 2023). Most of these accessions came from studies in *A. thaliana* with a few additional proteins identified in *Helianthus annuus*, *Lactuca sativa* and *Oryza sativa*. Details on original publications consulted for each accession can be found in the “Target capture design - loci overview” file (online supporting material).

#### c. Universal kits

To generate a direct comparison of intraspecific discriminatory power between genes from universal and custom target capture kits, the bait design for this study was expanded with loci commonly obtained with two universal kits. This comprised genes from the Angiosperms-353 (Johnson et al. 2019) and the Asteraceae Conserved Orthologous Sequence (Compositae-COS; Mandel et al. 2014) kits. The Angiosperms-353 kit captures single-copy nuclear genes that are conserved across flowering plants, based on large-scale transcriptome data (One Thousand Plant Transcriptomes Initiative 2019). Transcripts of single-copy nuclear genes from the Angiosperms-353 design were obtained from the closest relative to *Taraxacum* in this transcriptome dataset (*Tragopon porrifolius*, https://datacommons.cyverse.org/browse/iplant/home/shared/commons_repo/curated/oneKP_capstone_2019/transcript_assemblies/KGJF-Tragopogon_porrifolius) based on the list of ortholog names (https://github.com/smirarab/1kp/tree/master/alignments). To obtain orthologous sequences from the Compositae-COS kit, assembled exon sequences were obtained from the closest relative of *Taraxacum officinale* in the extensive phylogenomic dataset for Asteraceae (*Taraxacum kok-saghyz*; Mandel et al. 2019; https://doi.org/10.6084/m9.figshare.7697834). The obtained sequences from *Tragopogon porrifolius* (Angiosperms-353) and *Taraxacum kok-saghyz* (Compositae-COS) were translated into orthologous sequences from *Taraxacum officinale* following the pipeline described above. These exonic sequences are therefore comparable to what would normally be captured with the respective universal kits and, together with their intronic counterparts, were added to the bait design for this study. While the Angiosperms-353 kit was designed to have individual loci comprise full transcripts, loci from the Compositae-COS kit comprise individual exons, thereby affecting the amount of intronic sequence that could be included.

### 3. DNA isolation

Individuals belonging to the 17 accessions previously characterised with microsatellite data were grown from seed to obtain high-molecular-weight DNA samples. Seeds were germinated on wet filter paper in closed petri dishes (7 cm ø) for ten days (20 °C, 16 hr light/16 °C, 8 hr dark) in ECD01 incubators (Snijders Labs, Tilburg, The Netherlands). Seedlings were transferred to pots containing, volume-wise, 80% potting soil and 20% pumice, in a greenhouse. After two weeks, young leaves were sampled with an 8 mm hole-punching device – while avoiding the latex-rich leaf midrib – and fragments were flash-frozen in liquid nitrogen. DNA was isolated using the NucleoSpin Plant II kit (Macherey-Nagel, Dueren, Germany), following the manufacturer’s protocol.

For herbarium material, small leaf fragments (±1 cm^2^) were excised from the inner leaf parts of the specimen. As small historical samples yield very low DNA quantities, and are thus more vulnerable to contamination, historical DNA was isolated in a dedicated ultra-clean ancient-DNA laboratory facility at Naturalis Biodiversity Center. DNA was extracted using a modified CTAB protocol (Doyle and Doyle 1987), optimised for herbarium specimens with an isopropanol precipitation of 4 weeks (Quatela et al. 2023). DNA isolates were cleaned up with 0.9X NucleoMag (Machery-Nagel) magnetic beads, following recommendations by Quatela et al. (2023).

All DNA isolates were checked for purity and concentration using a Nanodrop device and Qubit 3^®^ fluorometer (HS kit, Thermo Fisher Scientific, Waltham, MA, USA), respectively. Historical DNA samples were checked for fragmentation patterns with high-sensitivity electrophoresis (Bioanalyzer 2100, Agilent, Santa Clara, CA, USA) to determine if, and to what extent, further fragmentation was necessary for short-read compatible library preparation. 11 herbarium DNA samples were not further fragmented (major genomic DNA peak <2000 bp). For the other samples, fragmentation to the appropriate size (300-350 bp) was then achieved through DNA shearing by sonication (Bioruptor^®^ Plus, Diagenode, Liège, Belgium) of 200 ng genomic DNA in 50 µL Tris-EDTA. The number of sonication cycles (30 seconds ON, 90 seconds OFF) was optimised by trialing two samples of different degradation with 6, 8, and 10 cycles and analysing the results with high-sensitivity electrophoresis (Bioanalyzer 2100, Agilent). As a result, five herbarium samples were subjected to 6 cycles (major peak in genomic DNA 2500-5000 bp) and two herbarium as well as all fresh samples to 8 cycles (major peak in gDNA >5000 bp).

For 11 accessions (6 fresh, 5 herbarium), a technical sequencing replicate was generated by taking a second amount of DNA from the initial extraction stock through fragmentation step into the library preparation, see library preparation sheet (online supporting material) for more information.

### 4. Library preparation, target capture & sequencing

DNA isolates were processed into libraries compatible with short-read Illumina^®^ sequencing using the NEBNext Ultra II library preparation kit for Illumina (New England Biolabs, Ipswich, MA, USA) with compatible dual-index barcodes for multiplexed sequencing (Ultra II primer set 1). Manufacturer protocols were followed with the exception that half-volumes of buffers and enzymes were deployed. Double-sided size selection was applied to all sonicated samples with NucleoMag beads (Machery-Nagel). PCR dual index barcoding and amplification was performed with 8 cycles following the set-up recommended in the NEBNext Ultra II protocol. Fragment distribution of all libraries was determined by high-sensitivity electrophoresis (Bioanalyzer 2100, Agilent) and concentration with a Spark^®^ 10M fluorescence plate reader (Tecan, Männendorf, Switzerland).

Libraries were subsequently divided into two equimolar pools based on average fragment size distribution: (1) 23 herbarium samples (230-368 bp) and (2) all 24 fresh samples + 1 herbarium sample (505-1157 bp). Using a SpeedVac vacuum concentrator (Thermo Fisher Scientific) the pools were spun dry and resuspended in 7µL Tris-EDTA. Libraries were enriched through hybridisation target capture with the custom designed *Taraxacum* kit (myBaits^®^ v5 chemistry, Daicel Arbor Biosciences). Manufacturer recommendations were followed with hybridisation at 65°C for 40 hours and a triple wash in 1.5 mL tubes. Post-capture amplification followed the recommendations in the myBaits v5 protocol with 14 cycles and using a KAPA HiFi HotStart ready master mix (Roche, Basel, Switzerland) and universal P5 and P7 primers (Thermo Fisher Scientific). Following clean-up using 0.9X NucleoMag beads, DNA concentration and fragment length distribution of the post-capture pools were quantified as above. Pools were finally combined in an equimolar pool (25 µL, 1.42 ng/µL) and sequenced on an Illumina^®^ NovaSeq with 150 bp paired end reads output (Novogene Europe, Cambridge, UK).

### 5. Bioinformatics

Sequence read processing and mapping followed recommendations by Gutiérrez-Valencia et al. (2022). Following raw read quality control using FastQC (Andrews 2010) and MultiQC (Ewels et al. 2016), raw reads were quality filtered and trimmed to remove adapters using Trimmomatic v0.39 (Bolger et al. 2014) with average phred-33 quality score cut-off of 28 and a minimum resulting sequence length of 40 nucleotides. Unpaired reads were combined into single files per sample for more streamlined processing. Reads were mapped separately for each category of loci against the relevant target capture reference sequence files (online supporting material) using BWA mem v0.7.17 (Li and Durbin 2009), quality-filtered (-q 20) processed into .bam files using samtools v1.6 (Danecek et al. 2021), and further filtered by removing duplicate reads using Picard v2.20.4 (https://broadinstitute.github.io/picard/).

Variants were called with FreeBayes v1.0.2 for triploid genomes. For nuclear genes (categories b and c), VCF files were split up using BCFtools v1.11 (Danecek et al. 2021) with the -R function using dedicated BED files (online supporting material). All VCF files were filtered using BCFtools to retain only high-quality and high-read-depth variants: variant call quality ≥20 (view -i ‘%QUAL>=20’); individual genotype quality ≥20 (filter -e ‘FMT/GQ<20’); read depth ≥15 (filter -e ‘FMT/DP<15’); with no missing genotypes (view - g ^miss) and no invariant sites (view -c 1:minor). To avoid double-counting variants spanning exon-intron boundaries, only single-nucleotide variants (SNVs) were retained to document genetic variation (view -i ‘strlen(REF)=1 && strlen(ALT)=1’).

To calculate the level of false positive variants captured in herbarium material, due to deamination bias (postmortem C → T transitions), which generally increases with age (Weiß et al. 2016), raw reads were investigated with mapDamage v2.2.3 (Jónson et al. 2013).

### 6. Analysis of Molecular Variance (AMOVA)

Analysis of molecular variance (AMOVA; Excoffier et al. 1992) was deployed to quantify differences in the distribution of the captured intraspecific variation between customised and universal target capture sequencing loci. AMOVA determines the distribution of intraspecific variation at different hierarchical levels: 1) between populations; 2) within populations; and 3) within individuals. Here, apomictic clonal lineages (ACLs) were defined as ‘populations’ to reveal what proportion of the captured intraspecific variation is discriminant on a larger scale (discrimination between ACLs) and a finer scale (discrimination between accessions within the same ACL). To quantify the effect of herbarium sampling on the captured intraspecific variation, a separate AMOVA was performed where source material (herbarium versus fresh) was analysed as an additional level of variation. In the latter exercise, samples from the ACL *obtusifrons* were omitted as this ACL was represented by only a single herbarium individual. For AMOVA, we used the option implemented in Poppr (Kamvar et al. 2014) that is based on the calculation of squared Euclidean distances among multilocus genotypes, which correctly reflects the clonal reproduction of the apomictic lineages. Therefore, it was not possible to determine within-individual variation. Thus, AMOVAs partitioned variation at two levels: between ACLs, and between individuals within the same ACL.

AMOVAs for polyploid data (Meirmans & Liu 2018) were performed for each locus category separately, using the R package Poppr v2.9.6 (Kamvar et al. 2014), which was specifically developed for analysing genotype data from (partially) clonal polyploid populations. The statistical support for each AMOVA was determined by making 1000 random permutations to the data matrix (e.g., randomising the identities of the samples) using the function ‘randtest’ of the R package ade4 v1.7-22 (Dray and Dafour 2007). The probability of the observed hierarchical structure in the genetic variants was determined as the proportion of random permutations that did not affect the structure.

For Custom genes (category b), each subcategory of functional genes (I-V) was analysed separately in an additional AMOVA, to determine the function of the most variable genes. For Universal genes (category c), the benefit of family-specific kit (Compositae-COS) compared to a broadly universal kit (Angiosperms-353) was tested in additional AMOVAs. Finally, separate AMOVAs were also performed for exons and introns for each (sub)category to reveal the differences in captured variation between coding and non-coding regions.

As the consistently large genetic distance between herbarium sample ‘H32’ and the other samples belonging to ACL *ekmanii* indicated a putatively misidentified sample, the effect of discarding this sample on AMOVA results was investigated for each category and subcategory of loci (supplementary material).

### 7. Principal Component Analysis (PCA)

To visualise the discriminatory power obtained with each category of loci, both between and within ACLs, the genetic variation was analysed in a principal component analysis (PCA). PCAs were performed in R with the function ‘dudi.pca’ from the package ade4 v1.7-22 (Dray and Dufour 2007) using 5 axes (nf=5), and visualised by plotting the coordinates of the two principal components capturing the most variation, colouring individual samples by ACL identity.

### 8. Genetic distance

To quantify the differentiation between individual members of the five ACLs, pairwise genetic distance matrices were calculated for each category of loci. Distances were calculated as Euclidean differences in allelic counts, using the R package adegenet v2.1.10 (Jombart 2008), which calculates the sum of squared allelic distances of all variants and takes the square root of this sum. In other words, the distance between genotypes 0/0/0 and 0/0/1 is √1^2^=1, but between genotypes 0/0/0 and 0/1/1 the distance is √(1^2^ + 1^2^)=√2≈1.41. This distance metric is the same one that underlies the calculation of the variance components in the AMOVA for polyploids (Meirmans and Liu 2018). Distance matrices were coloured according to a scale for each ACL individually.

To evaluate variation due to differences in capture efficacy, the distance between both technical replicates was calculated for the different categories of loci. This was contrasted with the distance between each technical replicate and the nearest other biological sample.

## Results

### 1. How reliable is target capture sequencing in herbarium specimens?

Sequencing of herbarium material produced on average 18.7 million reads per sample, of which 12.0% were filtered out and 14.2% were unpaired. Fresh material produced more reads with 22.7 million, fewer discarded in filtering (2.1%) and fewer unpaired (2.0%). The average on-target ratio (proportion of reads mapping to target) was higher for herbarium material (84.5%) than for fresh material (66.6%). This difference is explained by the greater proportion of reads mapping to GBS loci in herbarium samples (36.7%), about 10% higher than expected from bait design. In contrast, read depth was lower in herbarium samples (287, Fig. 1) compared to fresh samples (500), consistent with both the lower number of reads and the higher proportion lost in filtering. Read length was also shorter and more variable in herbarium samples (average 144 bp), whereas fresh samples consistently averaged 149 bp. Despite an age range spanning 46 years (1954-2000), there was no clear correlation between target DNA fragmentation and specimen age. There were also no signs of deamination bias (postmortem C → T transitions) in any of the herbarium material (more details in ‘mapDamage reports’, online supporting material), indicating high-quality DNA preservation in herbarium vouchers of *Taraxacum*, which are usually air dried upon collection.

One herbarium sample (H33, collected in 1983, belonging to ACL *obtusifrons*) clearly underperformed in all of these metrics, for both technical replicates. With fewer reads surviving filtering (<2.5 million) and a much lower read length (129-130 bp compared to the average 144 bp), the average read depth (36-41) was nearly half that of the other samples (≥78). As this drastically affected the number variants surviving filtering (due to the condition that no missing data is allowed for AMOVA in Poppr), this sample was not considered for downstream analysis.

Although enrichment success (percentage on-target reads) was comparable between technical replicates of the same sample, the resulting read depth varied considerably in 4 out of the 10 samples (H40, N22, N24, and N26) reflecting large differences in sequencing output. One technical replicate of an herbarium sample (rep_H36) failed as it had only 48,779 filtered reads and an average read length of 125 bp. This sample was therefore excluded from further analysis. More details on target enrichment statistics are available in the ‘Sequence-Mapping_results’ file in the online supporting material.

### 2. How many intraspecific variants can customised and universal kits yield?

The two customised target capture approaches yielded considerably more variants overall, but confidence varied due to uneven read depths. GBS loci produced the highest number of single-nucleotide variants (22,045, Table 1), yet only 8,927 SNVs (1.88% of the total length) passed filtering, corresponding to a loss of ∼60% (Fig. 1). Custom genes (exons and introns combined) were more consistent, with 14,962 SNVs (2.56% of the total length) retained after filtering from an original 20,194 SNVs, a loss of ∼26%. Universal genes showed a loss of ∼31%, resulting in 5,723 high-confidence SNVs, with a comparable proportion of variable sites (2.05% of the total length).

**Table 1:** Number of variants for each category of loci at different steps of the variant filtering process. Variants are categorised into single-(SNV) and multi-nucleotide (MNV, stretches of ≥2 nucleotides) variants, and insertions or deletions (indel). Multi-allelic variants (>2 alleles) are split up into separate variants. For SNVs, the percentage of variable positions in the target sequences is reported. Variant calling was based on read mapping to full genes, therefore there is overlap in multi-nucleotide variants and indels between exons and introns as some of these span the exon-intron boundaries. The three main approaches of target capture sequencing that are contrasted in this study are highlighted in bold.

| Locus category | Target length (bp) | Unfiltered dataset |  |  |  | Quality filtering* |  |  |
| --- | --- | --- | --- | --- | --- | --- | --- | --- |
|  |  | total | SNV | MNV | indel | SNV | MNV | indel |
| **GBS** | **460 853** | **35 974** | **22 045**<br>(4.66%) | **10 255** | **3 674** | **8 927**<br>(1.88%) | **6 337** | **1 777** |
| **Custom genes** | **571 911** | **35 038** | **20 194**<br>(3.45%) | **10 195** | **4 649** | **14 962**<br>(2.56%) | **8 089** | **3 863** |
| - exons | 334 174 | 18 175 | 11 230<br>(3.29%) | 4 955 | 1 990 | 8 615<br>(2.52%) | 4 298 | 1 734 |
| - introns | 212 350 | 15 222 | 7 893<br>(3.64%) | 4 715 | 2 614 | 5 587<br>(2.58%) | 3 387 | 2 107 |
| Universal genes | 426 070 | 22 840 | 13 990<br>(3.22%) | 6 015 | 2 835 | 10 022<br>(2.30%) | 4 748 | 2 226 |
| **- exons** | **273 117** | **12 523** | **8 361**<br>(3.01%) | **3 045** | **1 117** | **5 723**<br>(2.05%) | **2 341** | **837** |
| - introns | 152 953 | 10 453 | 5 629<br>(3.60%) | 3 052 | 1 772 | 4 299<br>(2.75%) | 2 472 | 1 436 |
| *Angiosperms-353* | 241 162 | 14 241 | 8 358<br>(3.40%) | 3 932 | 1 951 | 6 348<br>(2.58%) | 3 298 | 1 598 |
| - exons | 95 382 | 4 693 | 3 122<br>(3.22%) | 1 211 | 360 | 2 187<br>(2.25%) | 1 005 | 277 |
| - introns | 145 781 | 9 665 | 5 236<br>(3.52%) | 2 797 | 1 632 | 4 161<br>(2.79%) | 2 357 | 1 360 |
| *Compositae-COS* | 184 907 | 8 599 | 5 632<br>(2.99%) | 2 083 | 884 | 3 674<br>(1.95%) | 1 450 | 628 |
| - exons | 177 735 | 7 830 | 5 239<br>(2.89%) | 1 834 | 757 | 3 536<br>(1.95%) | 1 336 | 560 |
| - introns | 7 172 | 788 | 393<br>(5.37%) | 255 | 140 | 138<br>(1.91%) | 115 | 76 |

\*: Quality filtering criteria (bcftools): variant quality (QUAL $\geq 20$ ), genotype quality (GQ $\geq 20$ ), read depth (DP $\geq 15$ ), no missing genotypes (^miss).

Within the customised gene set, loci associated with heat resistance and pollen development were the most variable, at 2.73% and 2.66% of the total length, respectively (Table S1, supplementary material). Among the universal kits, exons from the family-specific Compositae-COS kit yielded more variants (3,536 after filtering, Table 1) than the broadly universal Angiosperms-353 kit (2,187), mainly due to the higher number of loci obtained with the Compositae-COS in *T. officinale* (442 compared to 193). However, proportionally speaking the loci from the Angiosperms-353 kit were more variable (2.25% of the total length compared to 1.95%).

Although the analysis presented here only focuses on SNVs to enable comparisons between exons and introns (i.e. some longer variants span the exon-intron boundary), multi-nucleotide variants (MNVs) and insertions/deletions (indels) comprise a considerable amount of additional intraspecific variation: 33.2-42.4% of all variants. Intronic regions proportionally contained more indels (16.9-17.2% of all the variants) than exonic regions (8.9-10.9%), where they could potentially disrupt the reading frame of the amino acid translation.

### 3. How informative are intraspecific variants from customised and universal kits?

#### a. Differences between ACLs vs differences between members of the same ACL

Although GBS loci yielded fewer high-confidence SNVs than using full genes from the customised approach, a higher proportion (31.35%, Table 2) of the variation was assigned to the deepest level of intraspecific variation: differences between individuals belonging to the same apomictic clonal lineage (Fig. 1). Custom genes and exons from universal kits assigned similar proportions of variation to this level, at 20.44% and 18.87%, respectively. Within the universal datasets, Angiosperms-353 exons showed 20.75% of the variation at this level, whereas Compositae-COS exons showed 17.71% (Table 2).

**Table 2:**
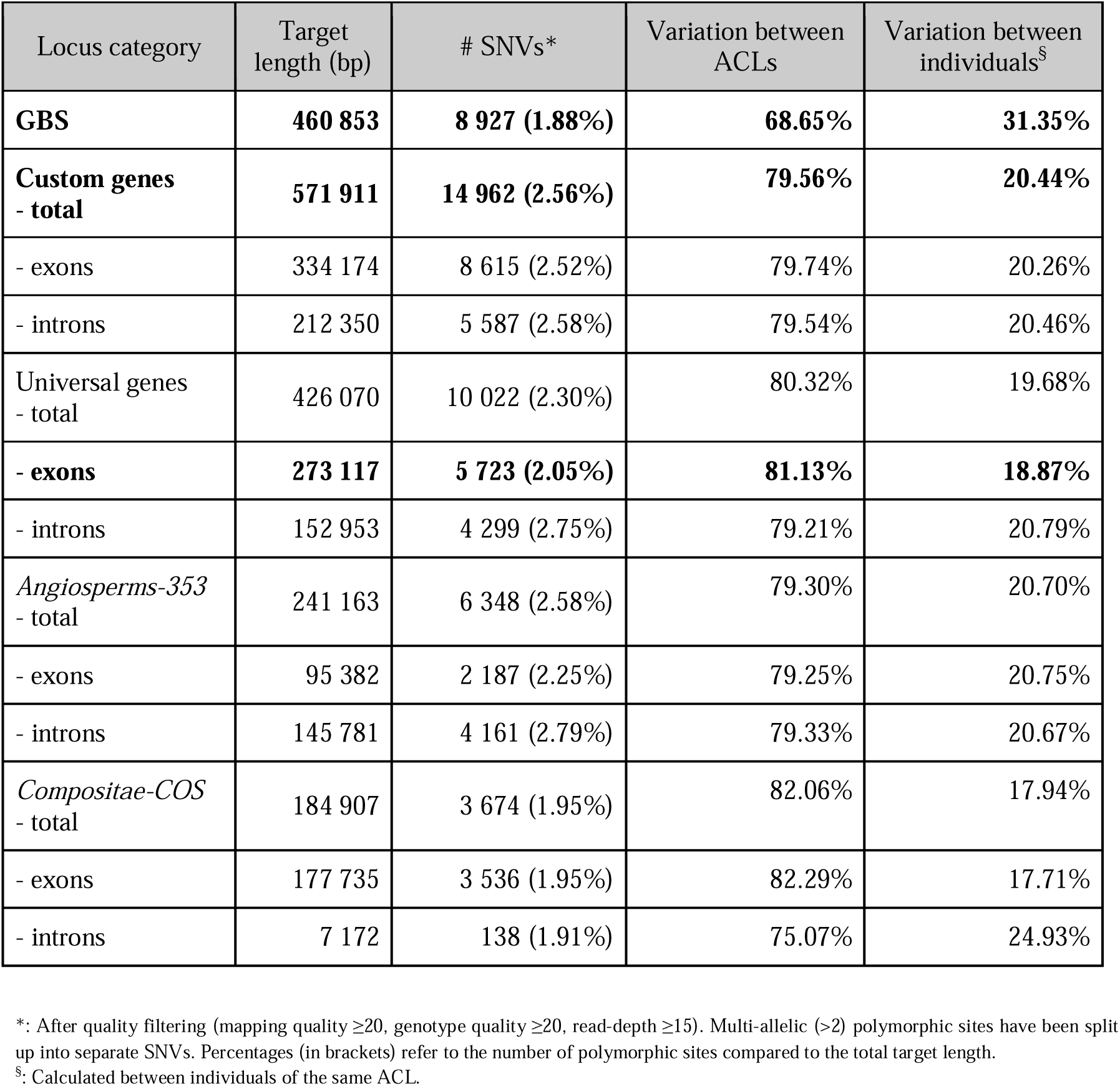
General AMOVA results for the comparison between GBS loci, custom selected genes, and genes from universal target capture kits. The observed genetic variation is characterised on two hierarchical levels: between and within apomictic clonal lineages (ACLs). Only single-nucleotide variants are considered to avoid double counting multi-nucleotide variants spanning exon-intron boundaries. The three main approaches of target capture sequencing that are contrasted in this study are highlighted in bold.

Differences between herbarium and fresh material of the same ACL were small, ranging from 1.25% in exons from universal kits to 3.34% using the customised GBS approach (Table 3). In fact, the variation between herbarium and fresh material captured with exons from universal kits had a statistically non-significant (p=0.112) structure. This was mainly caused by the extremely low level of separation (0.55% of the variation) at this level using exons from the Compositae-COS kit. Most of the variation within ACLs came from differences between samples of the same source, with one herbarium sample (H32 belonging to *ekmanii*) accounting for 4.4-5.2% of this variation (see reduced percentages from the AMOVA excluding sample H32 in Table S2, supplementary material).

**Table 3:** Three-level AMOVA results for the comparison between GBS loci, custom selected genes, and genes from universal target capture kits for four widespread apomictic clonal lineages (ACLs) of *Taraxacum officinale*. Compared to a two-level AMOVA, an extra hierarchical level of variation is considered in the form of herbarium versus fresh samples (called ‘source’ here) within ACLs. The variation between members of the same ACL is therefore divided into 1) variation between fresh and herbarium material (‘variation between sources’) and 2) variation between members of the same ACL of the same source material type (‘variation within sources’). As only one herbarium sample was analysed for *Obtusifrons*, all samples from this ACL were excluded. The three main approaches of target capture sequencing that are contrasted in this study are highlighted in bold.

| Locus category | # SNVs* | Variation between ACLs | Variation between sources | Variation within sources |
| --- | --- | --- | --- | --- |
| **GBS** | **7 199 (1.52%)** | **64.59%** | **3.34%** | **32.07%** |
| **Custom genes - total** | **13 644 (2.34%)** | **77.09%** | **2.24%** | **20.67%** |
| - exons | 7 859 (2.30%) | 77.39% | 2.10% | 20.51% |
| - introns | 5 108 (2.37%) | 76.90% | 2.39% | 20.71% |
| Universal genes - total | 8 894 (2.05%) | 78.13% | 1.51% | 20.36% |
| **- exons** | **4 990 (1.79%)** | **79.24%** | **1.25%^** | **19.52%** |
| - introns | 3 904 (2.50%) | 76.68% | 1.86% | 21.47% |
| *Angiosperms-353* - total | 5 715 (2.32%) | 76.77% | 2.02% | 21.20% |
| - exons | 1 933 (1.99%) | 76.90% | 2.35% | 20.75% |
| - introns | 3 782 (2.54%) | 76.71% | 1.85% | 21.44% |
| *Compositae-COS* - total | 3 179 (1.69%) | 80.54% | 0.60%^ | 18.86% |
| - exons | 3 057 (1.69%) | 80.71% | 0.55%^ | 18.74% |
| - introns | 122 (1.69%) | 75.53% | 1.97%^ | 22.51% |

\*: After quality filtering (mapping quality $\geq 20$ , genotype quality $\geq 20$ , read-depth $\geq 15$ ). Multi-allelic (>2 alleles) polymorphic sites have been split up into separate SNVs. Percentages (in brackets) refer to the number of polymorphic sites compared to the total target length.
^: No significant genetic structure at this level ( $p > 0.05$ after 1000 random permutations).

#### b. Separating near-identical genotypes of the same ACL

Relationships between and within apomictic clonal lineages (ACLs) of *T. officinale* were consistently recovered with both customised and universal approaches of target capture sequencing (Fig. 1, Fig. 3). All five ACLs formed distinguishable clusters in the principal component analysis (PCA), where three ACLs (*alatum*, *ekmanii*, and *interveniens*) contained one or more clearly distinct genotypes (Fig. 3). In *alatum*, two genotypes (fresh samples ‘N05’ and ‘N18’) were clearly distinct from the core of the ACL and from each other, a relationship that was equally well exposed in both customised and universal approaches. In *ekmanii*, one distinct genotype (fresh sample ‘N22’) was only visibly discriminated using the customised GBS approach (Fig. 3A). Herbarium samples generally clustered together with the core of the ACL, with the exception of herbarium sample ‘H32’ (belonging to *ekmanii*).

**Figure 3:**
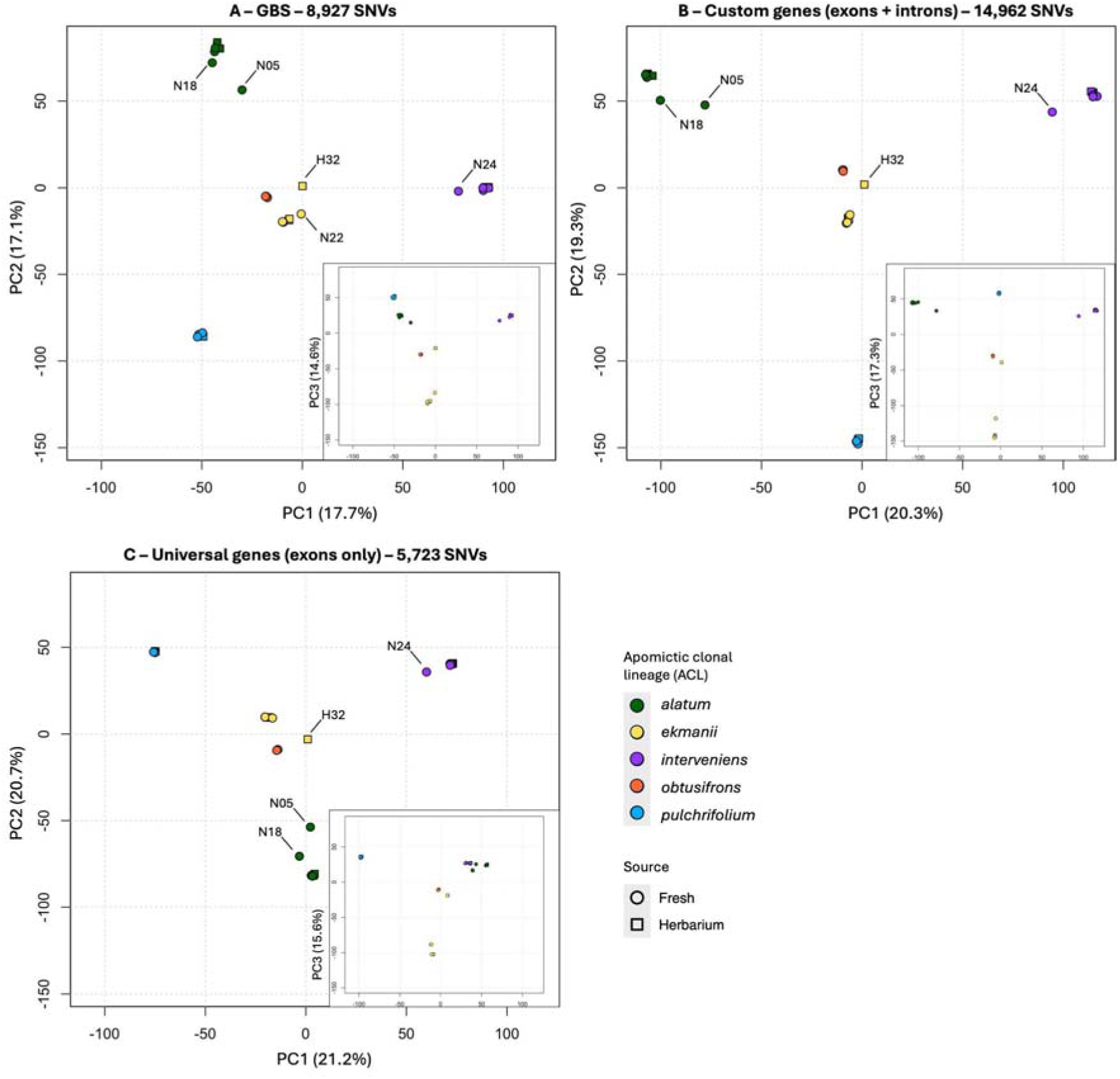
Principal component analysis (PCA) for different categories of loci. Comparison of apomictic clonal lineage discrimination through principal component analysis (PCA). Different categories of loci are contrasted to reveal the difference in discriminatory power between customised (A – GBS loci; B – Custom genes) and universal (C – Angiosperms-353 + Compositae-COS) target capture sequencing. Samples that can be discriminated from the core of the clonal lineage are highlighted, corresponding to sample numbering in Fig. 4 (Online Supporting Material - ‘Accession Information’ for more details). The small inserts comprise plots of the first and third principal components. Scales are identical in each plot. Conserved exons from universal target capture kits provided a discrimination comparable with custom genes and GBS loci, both between and within ACLs.

For these distinct genotypes within ACLs, the universal approach yielded genetic distances (Fig. 4) comparable to using the customised GBS approach. The customised full gene approach gave considerably higher genetic distances (33-47%) for these more distantly related samples (‘N05’ in *alatum*, ‘H32’ in *ekmanii*, and ‘N24’ in *interveniens*). However, when comparing the most closely related samples, separation was superior with the customised GBS approach (Fig. 1, Fig. 4). At this finest scale, the limitations of exons from universal kits are exposed, with distances about half the size achieved with customised approaches (i.e., d=14.6 compared to d=29.3-34.2 in *interveniens*). The lowest genetic distance was measured between two herbarium samples (‘H38’ and ‘H40’) belonging to the ACL *pulchrifolium*, going as low as d=13.5 with the universal approach. This corresponds to a difference of 13 variants with one alternative allele, or fewer variants if more than one allele is different.

**Figure 4:**
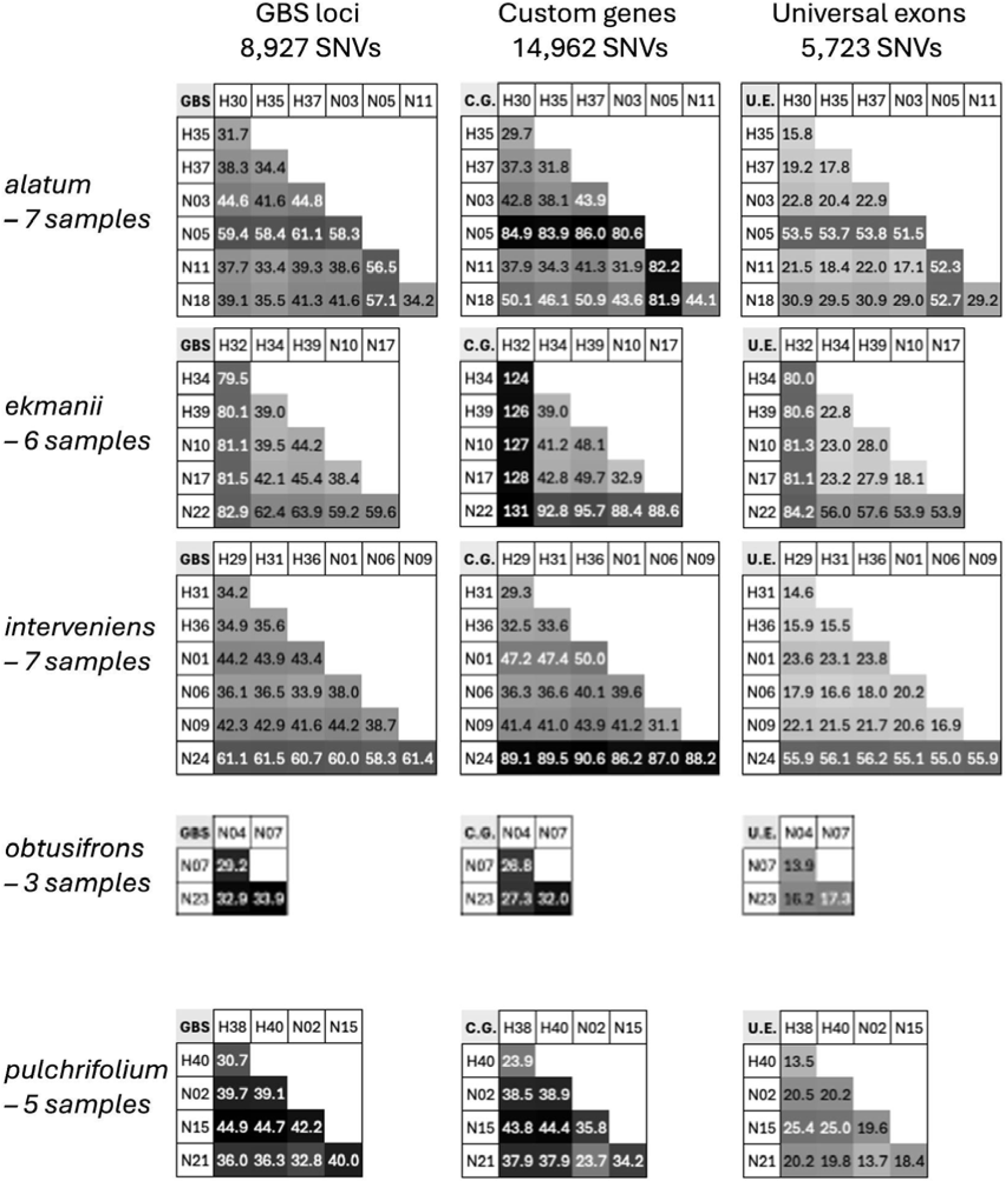
Heat maps of genetic distances between individual members within the five widespread apomictic clonal lineages (ACLs). Comparison is made between GBS loci, Custom genes (‘C.E.’; both exons and introns) and exons from universal kits (‘U.E.’; Angiosperms-353 and Compositae-COS). Distances were calculated as Euclidean distances based on allelic counts and are coloured according to a scale specific to the ACL. Distance values higher than the median for the respective ACL are indicated in white font. Details on individual accessions in the ‘Accession Information’ file (online supporting material).

In summary, customised approaches achieved superior discrimination between closely related samples, though distances were still measurable using universal exons. The customised GBS approach gave the finest resolution between near-identical genotypes, but was more sensitive to differences in read depth, as technical replicates could not always be distinguished from the closest biological relative (see ‘distance-replicates-check’ file in the online supporting material).

## Discussion

### 1. The power of universal target capture kits in intraspecific herbarium genomics

This study provides direct evidence for differences in genotyping accuracy between customised and universal target capture approaches, thereby giving meaningful answers to questions around the added value of (more costly and time-consuming) customised approaches (Fig. 1). The results will be highly relevant to the herbarium genomics community in choosing the most cost-effective approach for upscaling in herbarium-inclusive molecular ecology. Although customised approaches evidently capture more intraspecific variation and are thus better at discriminating closely related genotypes, a universal approach with exons from readily available kits recovered the same relationships between and within apomictic clonal lineages of dandelions. This highlights the utility of universal kits below the species level, with a high potential for upscaling due to the considerably lower cost and time invested.

Universal target capture kits are available for different taxonomic scales: for broad clades, such as all flowering plants (Angiosperms-353, Johnson et al. 2019), flagellate land plants (GoFlag451, Breinholt et al. 2021), or for specific plant families (i.e. Compositae-COS, Mandel et al. 2014; Fabaceae1005, Crameri et al. 2022). As this study was performed on dandelions (*Taraxacum officinale*, Compositae family), we targeted exons from both the Angiosperms-353 and the Compositae-COS universal kits. This dataset therefore also presents a direct comparison of genotyping precision below the species level between a broadly universal kit and a family-specific one. Interestingly, exons from the Angiosperms-353 kit yielded a higher proportion of variants: 2.25% single-nucleotide polymorphic sites compared to 1.95% for Compositae-COS. Nominally speaking, however, the Compositae-COS kit captured more variants as more targets were recovered in the *Taraxacum officinale* genome: 177,735 bp compared to 95,382 bp for Angiosperms-353, despite both kits having comparable target lengths (±260,000 bp). It is therefore advisable to use a family-specific kit when this is available, as more variants will be captured due to better target recovery.

Additional variation can be captured from the off-target reads: i.e., reads from nearby intronic and intergenic regions. Indeed, Yardeni et al. (2022) reported that >70% of the variation captured with Angiosperms-353 in samples of *Tillandsia* (Bromeliaceae) came from off-target regions. In the present study, intronic regions in loci from the Angiosperms-353 kit yielded nearly twice the number of variants located in the exonic counterparts (4,161 SNVs compared to 2,187, Fig. 1). Read depth will be considerably lower for these off-target regions, thereby affecting confidence in variant calling. However, variants from regions directly adjacent to the exons should be reliably captured, as they will be covered by overhanging parts of on-target reads (also called the ‘splash zone’; Woudstra et al. 2022). With updates to improve target recovery (mega353, McLay et al. 2021) and complementary versions of these kits adding more variable loci (Compositae-ParaLoss-1272, Moore-Pollard et al. 2024), universal target capture kits are becoming more reliable at capturing intraspecific variation.

### 2. The added value of customised target capture sequencing

If genomic resources are available, and budgetary constraints are less pressing, a customised approach can improve resolution considerably. GBS loci are more likely to be selectively neutral and are therefore preferred in assessing demography and population structure (Andrews et al. 2016). While these loci captured the most intraspecific variation overall, with 22,045 SNVs (4.66% of the total length, Table 1), variations in read depth make their capture less reliable. In the dataset generated here, only ±40% of the SNVs captured with GBS loci were recovered after filtering (Fig. 1). A possible explanation for this is a less efficient capture of these variable loci due to a higher mismatch between the oligonucleotide bait (a sequence from the *Taraxacum officinale* reference genome, Xiong et al. 2023) and the target fragments. Target capture baits can tolerate up to 30% mismatches, and capture efficiency drops off with increased genetic distance (Woudstra et al. 2022). The variants that do survive filtering, however, are generally more informative for this category of loci, with >31% of the variation attributed to differences between near-identical genotypes of the same apomictic clonal lineage (ACL). A higher intraspecific resolution is therefore achieved with a customised GBS approach.

Investing time in selecting variable genes with evolutionary adaptive potential is advised, particularly with the inclusion of introns, as these loci most reliably captured variation, confidently distinguishing between near-identical genotypes. The capture of functional genes allows the additional objective of revealing differences in environmental selection pressures, as highlighted in this study. In the complex of ACLs of the common dandelion (*Taraxacum officinale*) investigated here, the most variable genes were related to heat resistance and pollen development (Table S2, Supplementary Material). These ACLs are geographically widespread and have been found to adapt to urban heat islands (Woudstra et al. 2024), indicating potential underlying processes of diversifying selection pressures. Pollen development is likely affected by the transition to apomictic asexuality, which in *Taraxacum* happens completely independent of pollen (Van Dijk et al. 2009). With selection on healthy pollen production absent, the related genes are expected to accumulate mutations, eventually deteriorating to the point where pollen production is completely stopped (Meirmans et al. 2006). These examples highlight the evolutionary insights that can be gained with customised target capture in herbarium-inclusive molecular ecology.

### 3. Strategies for upscaling in herbarium-inclusive molecular ecology

Universal kits are considerably more affordable and accessible than customised target capture. Combining the broadly universal Angiosperms-353 kit with a family-specific kit, as done here, can yield more than 5700 SNVs far below the species level, at a combined cost (∼$266 per reaction, Table 4) below that of ordering a customised kit (∼$293). Considering the costs saved on computational resources and working hours a lot of money and time is saved. One target capture bait reaction can enrich up to 96 samples with high efficacy (Hale et al. 2020), reducing the cost to less than $3 per sample. Herbarium-inclusive population genomics and molecular ecology, sometimes requiring hundreds of samples, is therefore possible with universal kits, even under budgetary constraints.

**Table 4:**
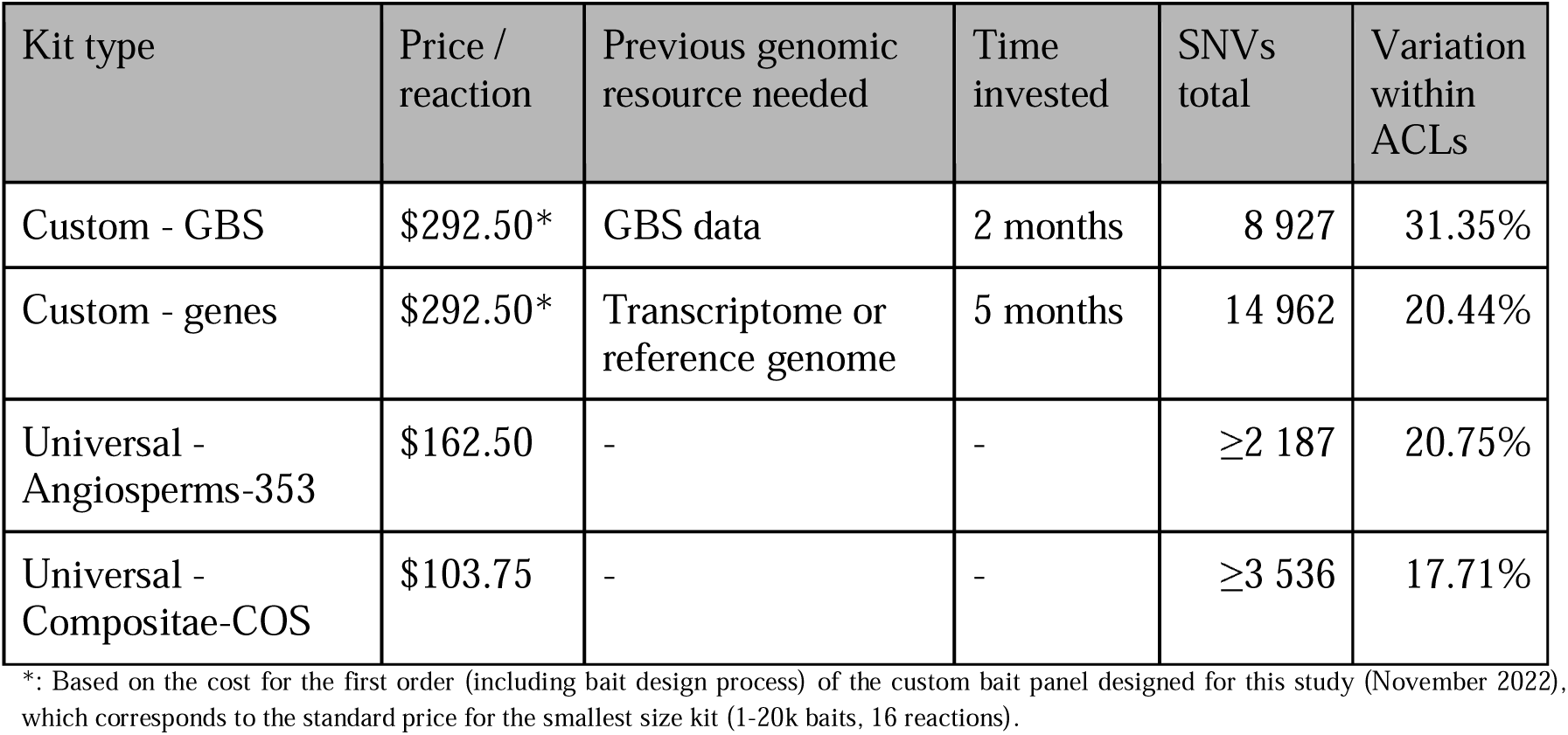
Return-on-investment comparison for different options of target capture sequencing kits. Prices are according to official listings by Daicel Arbor - myBaits^®^. Time invested in design is estimated as one researcher working full-time (40-hour work week), based on the design of the *Taraxacum* target capture kit designed for this study. Number of single-nucleotide variants (SNVs) is based on results after quality filtering for the five apomictic clonal lineages (ACLs) examined in this study, full details in Table 1 and 2.

| Kit type | Price / reaction | Previous genomic resource needed | Time invested | SNVs total | Variation within ACLs |
| --- | --- | --- | --- | --- | --- |
| Custom - GBS | \$292.50* | GBS data | 2 months | 8 927 | 31.35% |
| Custom - genes | \$292.50* | Transcriptome or reference genome | 5 months | 14 962 | 20.44% |
| Universal - Angiosperms-353 | \$162.50 | - | - | ≥2 187 | 20.75% |
| Universal - Compositae-COS | \$103.75 | - | - | ≥3 536 | 17.71% |

\*: Based on the cost for the first order (including bait design process) of the custom bait panel designed for this study (November 2022), which corresponds to the standard price for the smallest size kit (1-20k baits, 16 reactions).

An important factor is the quality of DNA isolates, for which herbarium-specific protocols now exist, making it possible to even extract long DNA fragments (>1 kbp) from herbarium material (Quatela et al. 2023). Indeed, DNA fragmentation by ultrasonication was still necessary for some of the herbarium samples in this study. Longer accurate reads (i.e. PacBio SMRT or Oxford Nanopore) would improve the possibility of phased read mapping and assembly, opening up the way for studying polyploid organisms (ubiquitous in the plant kingdom, Soltis et al. 2014), even from herbarium material. However, even heavily degraded DNA isolates, such as for ‘H37’ in this study with an average fragment length of 68 bp can be successfully enriched and sequenced (at an average read depth of 231). The ‘treasure chest’ represented by herbarium collections (Särkinen et al. 2012) can therefore be opened even further, below the species level. Universal target capture kits capture less intraspecific variation but allow more samples to be analysed at a lower cost. Therefore, less is arguably more in herbarium-inclusive molecular ecology.

## Acknowledgements

The authors thank Roxali Bijmoer, Laura van Hoek and Marcel Eurlings (Naturalis Biodiversity Center, Leiden) for help with access to the Oosterveld *Taraxacum* collection at the Dutch National Herbarium and the dedicated ancient DNA lab at Naturalis. We are grateful for the donation of verified seed material of apomictic clonal lineages of dandelion (*Taraxacum officinale*) by Jan Kirschner, Juhani Räsänen and Bob Trávníček. We additionally thank Jan Kirschner (Czech Academy of Sciences, Pruhonice), as well as Karst Meijer and Erik van den Ham (Herbarium Frisicum, Wolvega), for expert advice and verification of the identification of *Taraxacum* apomictic clonal lineages from herbarium material. We also thank Gregor Disveld (Netherlands Institute of Ecology, Wageningen) for help with the germination and propagation of dandelion material in the greenhouses.

Funding was provided by the Royal Netherlands Academy of Arts and Sciences (KNAW) in the form of an Institutenfonds grant to Koen Verhoeven and Barbara Gravendeel entitled “linking ecological analysis to plant and animal historical collections for tracking urban evolution”. Yannick Woudstra received a postdoctoral stipend from the Sven & Lily Lawski Foundation for Natural Sciences (grant N2023-0017) during his time at Stockholm University.

## Supplemental materials

A file with supplemental materials is available with this manuscript, comprising five tables and one figure referenced in the main text. The following contents are detailed in this supplemental file:

Table S1: Number of variants captured with different categories of Custom genes.

Table S2: Two-level analysis of molecular variance (AMOVA) for widespread apomictic clonal lineages, with putatively misidentified herbarium sample of *ekmanii* omitted.

Table S3: Two-level analysis of molecular variance (AMOVA) for widespread apomictic clonal lineages, all samples included.

Table S4: Two-level analysis of molecular variance (AMOVA) for widespread apomictic clonal lineages, focusing on different categories of genes with adaptive potential (Custom genes).

Table S5: Genetic distances between technical replicas of the same samples.

Figure S6: Principal component analysis (PCA) for different categories of loci, all samples included.

## Data availability statement

Raw sequence reads are deposited in the European Nucleotide Archive (ENA, hosted by EMBL-EBI) at BioProject PRJEB100583.

Other data and metadata related to this manuscript are deposited in the Zenodo repository at https://doi.org/10.5281/zenodo.17347176.

## Benefit sharing statement

A research collaboration was developed with scientists from the countries providing genetic samples (The Netherlands and Sweden) and all collaborators are included as co-authors.

Material from other countries (Czechia, Slovakia, Finland, Germany) was collected either from herbarium collections at Naturalis Biodiversity Center or donated to the senior author (Koen Verhoeven) by local collaborators (donations made before 12 October, which are acknowledged in the manuscript. The results of research have been shared with the provider communities and the broader scientific community (see Data availability statement above). More broadly, our group is committed to international scientific partnerships, as well as institutional capacity building.

## Author contributions

Yannick Woudstra, Koen Verhoeven, and Barbara Gravendeel conceptualised the study; Yannick Woudstra designed the target capture tool; Yannick Woudstra, Slavica Ivanovic and Niels Wagemaker performed the molecular lab work; Yannick Woudstra, Anne-Sophie Quatela, and Patrick Meirmans performed the bioinformatic analysis with input from Tanja Slotte; Yannick Woudstra led the writing of the manuscript. All authors contributed to the critical revision of the manuscript and approved the publication.

## References

Altschul, S. F., Gish, W., Miller, W., Myers, E. W., and Lipman, D. J. 1990. “Basic local alignment search tool”. Journal of Molecular Biology 215: 403–410.

Andrews, K. R., Good, J. M., Miller, M. R., Luikart, G., and Hohenlohe, P. A. 2016. “Harnessing the power of RADseq for ecological and evolutionary genomics”. Nature Reviews Genetics 17: 81–92. 10.1038/nrg.2015.28.

Andrews, S. 2010. “FastQC: A Quality Control Tool for High Throughput Sequence Data”. Available online at: http://www.bioinformatics.babraham.ac.uk/projects/fastqc/.

Bakker, F. T., Bieker, V., Martin, M. D. 2020. “Editorial: Herbarium Collection-Based Plant Evolutionary Genetics and Genomics”. Frontiers in Ecology and Evolution: 10.3389/fevo.2020.603948.

Barreiro, F. S., Vieira, F. G., Martin, M. D., Haile, J., Gilbert, T. P., Wales, N. 2017. “Characterizing restriction enzyme-associated loci in historic ragweed (*Ambrosia artemisiifolia*) voucher specimens using custom-designed RNA probes”. Molecular Ecology Resources 17(2): 209–220. 10.1111/1755-0998.12610.

Beck, J. B., Markley, M. L., Zielke, M. G., Thomas, J. R., Hale, H. J., Williams, L. D., and Johnson, M. G. 2021. “Are Palmer’s Elm-Leaf Goldenrod and the Smooth Elm-Leaf Goldenrod Real? The Angiosperms353 Kit Provides Within-Species Signal in *Solidago ulmifolia* s. l.” Systematic Botany 46(4): 1107–1113. 10.1600/036364421X16370109698740.

Bolger, A. M., Lohse, M., and Usadel, B. 2014. “Trimmomatic: a flexible trimmer for Illumina sequence data”. Bioinformatics 30(15): 2114–2120. 10.1093/bioinformatics/btu170.

Bouché, F., Lobet, G., Tocquin, P., and Périlleux, C. 2015. “FLOR-ID: an interactive database of flowering-time gene networks in *Arabidopsis thaliana*.” Nucleic Acids Research 44(D1): D1167–D1171. 10.1093/nar/gkv1054.

Breinholt, J. W., Carey, S. B., Tiley, G. P. et al. 2021. “A target enrichment probe set for resolving the flagellate land plant tree of life”. Applications in Plant Sciences 9(1): e11406. 10.1002/aps3.11406.

Crameri, S., Fior, S., Zoller, S., and Widmer, A. 2022. “A target capture approach for phylogenomic analyses at multiple evolutionary timescales in rosewoods (*Dalbergia* spp.) and the legume family (Fabaceae)”. Molecular Ecology Resources 22(8): 3087–3105. 10.1111/1755-0998.13666.

Danecek, P., Bonfield, J. K., Liddle, J., et al. 2021. “Twelve years of SAMtools and BCFtools”. GigaScience 10(2): giab008. 10.1093/gigascience/giab008.

Dang, Z., Li, J., Liu, Y., et al. 2024. “RADseq-based population genomic analysis and environmental adaptation of rare and endangered recretohalophyte *Reaumuria trigyna*”. The Plant Genome 17(1): e20303. 10.1002/tpg2.20303.

Dodsworth, S., Pokorny, L., Johnson, M. G., et al. 2019. “Hyb-Seq for Flowering Plant Systematics”. Trends in Plant Science 24(10): 887–891. 10.1016/j.tplants.2019.07.011.

Doyle, J. J., and Doyle, J. L. 1987. “A Rapid DNA Isolation Procedure for Small Quantities of Fresh Leaf Tissue”. Phytochemical Bulletin 19: 11–15.

Dray, S. and Dufour, A.-B. 2007. “The ade4 Package: Implementing the Duality Diagram for Ecologists”. Journal of Statistical Software 22(4): 1–20. 10.18637/jss.v022.i04.

Elshire, R. J., Claubitz, J. C., Sun, Q., Poland, J. A., Kawamoto, K., Buckler, E. S., and Mitchell, S. E. 2011. “A Robust, Simple Genotyping-by-Sequencing (GBS) Approach for High Diversity Species”. PLOS ONE 6(5): e19379. 10.1371/journal.pone.0019379.

Ewels, P., Magnusson, M., Lundin, S., and Käller, M. 2016. “MultiQC: summarize analysis results for multiple tools and samples in a single report”. Bioinformatics 32(19): 3047–3048. 10.1093/bioinformatics/btw354.

Excoffier, L., Smouse, P. E., and Quattro, J. M. 1992. “Analysis of molecular variance inferred from metric distances among DNA haplotypes: application to human mitochondrial DNA restriction data.” Genetics 131(2): 479–491. 10.1093/genetics/131.2.479.

Gahwens, F., Postuma, M., van Antro, M., et al. 2022. “epiGBS2: Improvements and evaluation of highly multiplexed, epiGBS-based reduced representation bisulfite sequencing.” Molecular Ecology Resources 22(5): 2087–2104. 10.1111/1755-0998.13597.

Gutiérrez-Valencia, J., Fracassetti, M., Berdan, E. L., et al. 2022. “Genomic analyses of the Linum distyly supergene reveal convergent evolution at the molecular level”. Current Biology 32(20): P4360–4371.e6. 10.1016/j.cub.2022.08.042.

Hale, H., Gardner, E. M., Viruel, J., Pokorny, L., and Johnson, M. G. 2020. “Strategies for reducing per-sample costs in target capture sequencing for phylogenomics and population genomics in plants”. Applications in Plant Science 8(4): e11337. 10.1002/aps3.11337.

Ibañez, V. N., van Antro, M., Peña-Ponton, C., Milanovic-Ivanovic, S., Wagemaker, C. A. M., Gawehns, F., and Verhoeven, K. J. F. 2023. “Environmental and genealogical effects on DNA methylation in a widespread apomictic dandelion lineage.” Journal of Evolutionary Biology 36(4): 663–674. 10.1111/jeb.14162.

Johnson, M. G., Pokorny, L., Dodsworth, S., et al. 2019. “A Universal Probe Set for Targeted Sequencing of 353 Nuclear Genes from Any Flowering Plant Designed Using k-Medoids Clustering”. Systematic Biology 68(4): 594–606. 10.1093/sysbio/syy086.

Jónson, H., Ginolhac, A., Schubert, M., Johnson, P. L. F., and Orlando, L. 2013. “mapDamage2.0: fast approximate Bayesian estimates of ancient DNA damage parameters”. Bioinformatics 29(13): 1682–1684. 10.1093/bioinformatics/btt193.

Jombart, T. 2008. “adegenet: a R package for the multivariate analysis of genetic markers.” Bioinformatics 24(11): 1403–1405. 10.1093/bioinformatics/btn129.

Kamvar, Z. N., Tabima, J. F., and Grünwald, N. J. 2014. “Poppr: an R package for genetic analysis of populations with clonal, partially clonal, and/or sexual reproduction.” PeerJ 2: e281. 10.7717/peerj.281.

Kirschner, J., and Štěpánek, J. 1996. “Modes of speciation and evolution of the sections in Taraxacum”. Folia Geobotanica 31: 415–426. 10.1007/BF02815386.

Kirschner, J., Oplaat, C., Verhoeven, K. J. F., et al. 2016. “Identification of oligoclonal agamospermous microspecies: Taxonomic specialists versus microsatellites”. Preslia 88: 1–17. http://www.preslia.cz/P161Kirschner.pdf.

Lang, P. L. M., Weiß, C. L., Kersten, S., et al. 2020. “Hybridization ddRAD-sequencing for population genomics of nonmodel plants using highly degraded historical specimen DNA”. Molecular Ecology Resources 20(5): 1228–1247. 10.1111/1755-0998.13168.

Lang, P. L. M., Erberich, J. M., Lopez, L, et al. 2024. “Century-long timelines of herbarium genomes predict plant stomatal response to climate change”. Nature Ecology & Evolution 8: 1641–1653. 10.1038/s41559-024-02481-x.

Liang, K. J., Shepher-Clowes, A., Agut, A., Hidalgo, O., Tejero Ibarra, P., and Viruel, J. 2025. “Conservation Implications for the Iberian Narrow Endemic *Androsace cantabrica* (Primulaceae) Using Population Genomics With Target Capture Sequence Data”. Ecology and Evolution 15(8): e71901. 10.1002/ece3.71901.

Li, H., and Durbin, R. 2009. “Fast and accurate short read alignment with Burrows-Wheeler Transform”. Bioinformatics, 25: 1754–1760. 10.1093/bioinformatics/btp324.

Mandel, J. R., Dikow, R. B., Funk, V. A., et al. 2014. “A target enrichment method for gathering phylogenetic information from hundreds of loci: An example from the Compositae” Applications in Plant Sciences 2(2): 1300085. 10.3732/apps.1300085.

Mandel, J. R., Dikow, R. B., Siniscalchi, C. M., et al. 2019. “A fully resolved backbone phylogeny reveals numerous dispersals and explosive diversifications throughout the history of Asteraceae”. Proceedings of the National Academy of Sciences of the United States of America 116(28): 14083–14088. 10.1073/pnas.1903871116.

Manzanilla, V., Teixidor-Toneu, I., Martin, G. J., Hollingsworth, P. M., De Boer, H. J., and Kool, A. 2022. “Using target capture to address conservation challenges: Population-level tracking of a globally-traded herbal medicine”. Molecular Ecology Resources 22(1): 212–224. 10.1111/1755-0998.13472.

McLay, T. G. B., Birch, J. L., Gunn, B. F. et al. 2021. “New targets acquired: Improving locus recovery from the Angiosperms353 probe set”. Applications in Plant Sciences 9(7): e11420. 10.1002/aps3.11420.

Meirmans, P. G., Den Nijs, H. C. M., and Van Tienderen, P. H. 2006. “Male sterility in triploid dandelions: asexual females vs asexual hermaphrodites”. Heredity 96: 45–52. 10.1038/sj.hdy.6800750.

Meirmans, P. G., and Liu, S. 2018. “Analysis of Molecular Variance (AMOVA) for Autopolyploids”. Frontiers in Ecology and Evolution: 10.3389/fevo.2018.00066

Michel, T., Tseng, Y.-H., Wilson, H., Chung, K.-F., Thomas, D. C., and Kidner, C. 2022. “A Hybrid Capture Bait Set for *Begonia*”. Edinburgh Journal of Botany 79: 1–33. 10.24823/ejb.2022.409.

Moore-Pollard, E. R., Jones, D. S., and Mandel, J. R. 2024. “Compositae-ParaLoss-1272: A complementary sunflower-specific probe set reduces paralogs in phylogenomic analyses of complex systems”. Applications in Plant Sciences 12(1): e11568. 10.1002/aps3.11568.

Narum, S. R., Buerkle, C. A., Davey, J. W., Miller, M. R., and Hohenlohe, P. A. 2013. “Genotyping-by-sequencing in ecological and conservation genomics”. Molecular Ecology 22(11): 2841–2847. 10.1111/mec.12350.

One Thousand Plant Transcriptomes Initiative: Leebens-Mack, J. H., Barker, M. S., Carpenter, E. J. et al. 2019. “One thousand plant transcriptomes and the phylogenomics of green plants”. Nature 547: 679–685. 10.1038/s41586-019-1693-2

Otero, E. P., Pimentel, M., Balbuena, E. S., and Piñeiro, R. 2024. “Phylogenomic support for the allopolyploid origin of the northwest Iberian endemic orchid Dactylorhiza cantabrica with Hyb-Seq”. Journal of Systematics and Evolution: 10.1111/jse.13131.

Ousmael, K. M., and Hansen, O. K. 2025. “From phylogenomics to breeding: Can universal target capture probes be used in the development of SNP markers for kinship analysis?” Applications in Plant Sciences 13(1): e11624. 10.1002/aps3.11624.

Pan, J., Wang, B., Pei, Z.-Y., Zhao, W., Gao, J., Mao, J.-F., and Wang, X.-R. 2015. “Optimization of the genotyping-by-sequencing strategy for population genomic analysis in conifers. Moleculary Ecology Resources 15(4): 711–722. 10.1111/1755-0998.12342.

Quatela, A.-S., Cangren, P., Jafari, F., Michel, T., De Boer, H. J., and Oxelman, B. 2023. “Retrieval of long DNA reads from herbarium specimens”. AoBPlants 15(6): plad074. 10.1093/aobpla/plad074.

Quatela, A.-S., Cangren, P., de Lima Ferreira, P., Woudstra, Y., Zsoldos-Skahjem, A., Bacon, C. D., de Boer, H. J., and Oxelman, B. 2025. “Phylogenetic relationships and the identification of allopolyploidy in circumpolar *Silene* sect. *Physolychnis*.” American Journal of Botany 112(6): e70051. 10.1002/ajb2.70051.

Quinlan, A. R., and Hall, I. M. 2010. “BEDTools: a flexible suite of utilities for comparing genomic features.” Bioinformatics 26(6): 841–842.

Richards, A. J. 1996. “Genetic variability in obligate apomicts of the genus *Taraxacum*”. Folia Geobotanica 31: 405–414. 10.1007/BF02815385.

Rosche, C., Baasch, A., Runge, K., Brade, P., Träger, S., Parisod, C., and Hensen, I. 2022. “Tracking population genetic signatures of local extinction with herbarium specimens”. Annals of Botany 129(7): 857–868. 10.1093/aob/mcac061.

Särkinen, T., Staats, M., Richardson, J. E., Cowan, R. S., Bakker, F. T. 2012. “How to Open the Treasure Chest? Optimising DNA Extraction from Herbarium Specimens”. PLoS ONE 7(8): e43808. 10.1371/journal.pone.0043808.

Slimp, M., Williams, L. D., Hale, H., and Johnson, M. G. 2021. “On the potential of Angiosperms353 for population genomic studies”. Applications in Plant Sciences 9(7): e11419. 10.1002/aps3.11419.

Soltis, D. E., Visger, J. C., and Soltis, P. S. 2014. “The polyploidy revolution then…and now: Stebbins revisited”. American Journal of Botany 101(7): 1057–1078. 10.3732/ajb.1400178.

Staats, M., Cuenca, A., Richardson, J. E., Vrielink-van Ginkel, R., Petersen, G., Seberg, O., and Bakker, F. T. 2011. “DNA Damage in Plant Herbarium Tissue”. PLOS ONE 6(12): e28448. 10.1371/journal.pone.0028448.

Underwood, C. J., Vijverberg, K., Rigola, D. et al. 2022. “A PARTHENOGENESIS allele from apomictic dandelion can induce egg cell division without fertilization in lettuce.” Nature Genetics 54: 84–93. 10.1038/s41588-021-00984-y.

Ter Hoeven, N. 2015. “blast2bed” (python script): https://github.com/nterhoeven/blast2bed. Accessed on 2022-07-15.

Van Dijk, P. J., and Bakx-Schotman, J. M. T. 2004. “Formation of Unreduced Megaspores (Diplospory) in Apomictic Dandelions (*Taraxacum officinale*, s.l.) Is Controlled by a Sex-Specific Dominant Locus.” Genetics 166(1): 483–492. 10.1534/genetics.166.1.483.

Van Dijk, P. J., de Jong, H., Vijverberg, K., and Biere, A. 2009. “An Apomixis-Gene’s View on Dandelions”. In: Schön, I., Martens, K., van Dijk, P. J. (eds), Lost Sex. Springer, Dordrecht. 10.1007/978-90-481-2770-2_22.

Veltman, M. A., Anthoons, B., Schrøder-Nielsen, A., Gravendeel, B., and De Boer, H. J. 2024. “Orchidinae-205: A new genome-wide custom bait set for studying the evolution, systematics, and trade of terrestrial orchids”. Molecular Ecology Resources 24(6): e13986. 10.1111/1755-0998.13986.

Weiß, C. L., Schuenemann, V. J., Devos, J., et al. 2016. “Temporal patterns of damage and decay kinetics of DNA retrieved from plant herbarium specimens”. Royal Society Open Science 3(6): 160239. 10.1098/rsos.160239.

Woudstra, Y., Kraaiveld, R., Jorritsma, A., et al. 2024. “Some like it hot: adaptation to the urban heat island in common dandelion”. Evolution Letters 8(6): 881–892. 10.1093/evlett/qrae040.

Woudstra, Y., Quatela, A.-S., Kidner, C., et al. 2022. “Chapter 14 Target Capture”. In: De Boer, H. J., Rydmark, M. O., Verstraete, B., and Gravendeel, B. (eds.), Molecular Identification of Plants: From Sequence to Species. Advanced Books: 10.3897/ab.e98875.

Woudstra, Y., Rees, P., Rakotoarisoa, S. E., Howard, C., Rønsted, N., and Grace, O. M. 2024. “An updated DNA barcoding tool for Aloe vera and related CITES-regulated species”. Conservation Letters 18(4): e13127. 10.1111/conl.13127.

Woudstra, Y., Viruel, J., Fritzsche, M., Bleazard, T., Mate, R., Howard, C., Rønsted, N., and Grace, O. M. 2021. “A customised target capture sequencing tool for molecular identification of *Aloe vera* and relatives”. Scientific Reports 11: 24347. 10.1038/s41598-021-03300-0.

Xiong, W., Risse, J., Berke, L. et al. 2023. “Phylogenomic analysis provides insights into MADS-box and TCP gene diversification and floral development of the Asteraceae, supported by de novo genome and transcriptome sequences from dandelion (*Taraxacum officinale*)”. Frontiers in Plant Science: 10.3389/fpls.2023.1198909.

Yardeni, G., Viruel, J., Paris, M., et al. 2022. “Taxon-specific or universal? Using target capture to study the evolutionary history of rapid radiations”. Molecular Ecology Resources 22(3): 927–945. 10.1111/1755-0998.13523.

