## Supplemental materials for "“Universal Hyb-Seq kits capture considerable intraspecific variation: Less is more in herbarium-inclusive molecular ecology”"

Contents:

Table S1: Number of variants captured with different categories of Custom genes.
Table S2: Two-level analysis of molecular variance (AMOVA) for widespread apomictic clonal lineages, with putatively misidentified herbarium sample of *ekmanii* omitted.
Table S3: Two-level analysis of molecular variance (AMOVA) for widespread apomictic clonal lineages, all samples included.
Table S4: Two-level analysis of molecular variance (AMOVA) for widespread apomictic clonal lineages, focusing on different categories of genes with adaptive potential (Custom genes).
Table S5: Genetic distances between technical replicas of the same samples.
Figure S6: Principal component analysis (PCA) for different categories of loci, all samples included.

Table S1: Number of variants captured with different categories of Custom genes, at different stages of the filtering process. Variants are categorised into single- (SNV) and multi-nucleotide (MNV, stretches of ≥2 nucleotides) variants, and insertions or deletions (indel). Multi-allelic variants (>2 alleles) are split up into separate variants. For SNVs, the percentage of variable positions in the target sequences is reported. Variant calling was based on read mapping to full genes, therefore there is overlap in multi-nucleotide variants and indels between exons and introns as some of these span the exon-intron boundaries.

| Locus category | Target length (bp) | Unfiltered dataset | | | | Quality filtering* | | |
|---|---|---|---|---|---|---|---|---|
| | | total | SNV | MNV | indel | SNV | MNV | indel |
| Apomixis^ | 25 387 | 1 986 | 1 071 (4.11%) | 673 | 242 | 760 (2.91%) | 539 | 193 |
| Flowering time regulation | 120 666 | 7 178 | 4 226 (3.42%) | 2 054 | 898 | 2 889 (2.34%) | 1 437 | 740 |
| - exons | 66 638 | 3 460 | 2 172 (3.17%) | 925 | 363 | 1 639 (2.39%) | 823 | 324 |
| - introns | 54 028 | 3 791 | 2 054 (3.72%) | 1 154 | 583 | 1 250 (2.26%) | 638 | 456 |
| Heat resistance | 81 582 | 5 085 | 2 928 (3.52%) | 1 526 | 631 | 2 263 (2.73%) | 1 266 | 529 |
| - exons | 51 576 | 3 068 | 1 812 (3.44%) | 938 | 318 | 1 399 (2.66%) | 785 | 275 |
| - introns | 30 006 | 2 057 | 1 116 (3.67%) | 604 | 337 | 864 (2.84%) | 495 | 275 |
| Pollen development | 272 591 | 16 938 | 9 727 (3.49%) | 4 912 | 2 299 | 7 416 (2.66%) | 4 055 | 1 940 |
| - exons | 173 714 | 9 717 | 5 984 (3.37%) | 2 659 | 1 074 | 4 637 (2.60%) | 2 330 | 958 |
| - introns | 98 877 | 7 423 | 3 743 (3.69%) | 2 351 | 1 329 | 2 779 (2.75%) | 1 816 | 1 074 |
| Stomatal regulation | 71 685 | 3 851 | 2 242 (3.07%) | 1 030 | 579 | 1 634 (2.25%) | 792 | 461 |
| - exons | 42 246 | 1 930 | 1 262 (2.93%) | 433 | 235 | 940 (2.19%) | 360 | 177 |
| - introns | 29 439 | 1 951 | 980 (3.27%) | 606 | 365 | 694 (2.33%) | 438 | 302 |

Table S2: Two-level analysis of molecular variance (AMOVA) for widespread apomictic clonal lineages, with putatively misidentified herbarium sample of *Ekmanii* omitted. The observed genetic variation is characterised on two hierarchical levels: between and within apomictic clonal lineages (ACLs). Only single-nucleotide variants are considered to avoid double counting multi-nucleotide variants spanning exon-intron boundaries.

| Locus category | Target length (bp) | # SNVs* | Variation between ACLs | Variation between individuals§ |
|---|---|---|---|---|
| **GBS** | **460 853** | **8 651 (1.82%)** | **73.09%** | **26.91%** |
| **Custom genes - total** | **571 911** | **14 549 (2.49%)** | **84.50%** | **15.50%** |
| - exons | 334 174 | 8 361 (2.45%) | 84.81% | 15.19% |
| - introns | 212 350 | 5 439 (2.52%) | 84.35% | 15.65% |
| Universal genes - total | 426 070 | 9 713 (2.23%) | 85.59% | 14.41% |
| **- exons** | **273 117** | **5 540 (1.99%)** | **86.37%** | **13.63%** |
| - introns | 152 953 | 4 173 (2.67%) | 84.53% | 15.47% |
| *Angiosperms-353* - total | 241 163 | 6 163 (2.50%) | 84.65% | 15.35% |
| - exons | 95 382 | 2 127 (2.19%) | 84.57% | 15.43% |
| - introns | 145 781 | 4 036 (2.71%) | 84.69% | 15.31% |
| *Compositae-COS* - total | 184 907 | 3 550 (1.88%) | 87.20% | 12.80% |
| - exons | 177 735 | 3 413 (1.88%) | 87.47% | 12.53% |
| - introns | 7 172 | 137 (1.90%) | 78.77% | 21.23% |

*: After quality filtering (mapping quality ≥20, genotype quality ≥20, read-depth ≥15). Multi-allelic (>2) polymorphic sites have been split up into separate SNVs. Percentages (in brackets) refer to the number of polymorphic sites compared to the total target length.
§: Calculated between individuals of the same ACL.

Table S3: Two-level analysis of molecular variance (AMOVA) for widespread apomictic clonal lineages, all samples included. The observed genetic variation is characterised on two hierarchical levels: between and within apomictic clonal lineages (ACLs). Only single-nucleotide variants are considered to avoid double counting multi-nucleotide variants spanning exon-intron boundaries.

| Locus category | Target length (bp) | # SNVs* | Variation between ACLs | Variation between individuals§ |
|---|---|---|---|---|
| **GBS** | **460 853** | 5 893 (1.19%) | 63.34% | 36.66% |
| **Custom genes - total** | **571 911** | 10 748 (1.84%) | 78.91% | 21.09% |
| - exons | 334 174 | 6 528 (1.91%) | 79.32% | 20.68% |
| - introns | 212 350 | 3 611 (1.67%) | 78.43% | 21.57% |
| Universal genes - total | 426 070 | 6 866 (1.58%) | 80.09% | 19.91% |
| **- exons** | **273 117** | 4 115 (1.48%) | 81.00% | 19.00% |
| - introns | 152 953 | 2 751 (1.76%) | 78.66% | 21.34% |
| *Angiosperms-353* - total | 241 163 | 4 200 (1.70%) | 78.86% | 21.14% |
| - exons | 95 382 | 1 514 (1.56%) | 79.09% | 20.91% |
| - introns | 145 781 | 2 686 (1.80%) | 78.72% | 21.28% |
| *Compositae-COS* - total | 184 907 | 2 666 (1.41%) | 81.97% | 18.03% |
| - exons | 177 735 | 2 601 (1.43%) | 82.10% | 17.90% |
| - introns | 7 172 | 65 (0.91%) | 75.40% | 24.60% |

*: After quality filtering (mapping quality ≥20, genotype quality ≥20, read-depth ≥15). Multi-allelic (>2) polymorphic sites have been split up into separate SNVs. Percentages (in brackets) refer to the number of polymorphic sites compared to the total target length.
§: Calculated between individuals of the same ACL.

Table S4: Two-level analysis of molecular variance (AMOVA) for widespread apomictic clonal lineages, focusing on different categories of genes with adaptive potential (Custom genes). The observed genetic variation is characterised on two hierarchical levels: between and within apomictic clonal lineages (ACLs). Only single-nucleotide variants are considered to avoid double counting multi-nucleotide variants spanning exon-intron boundaries.

| Locus category | Target length (bp) | # SNVs* | Variation between ACLs | Variation between individuals§ |
|---|---|---|---|---|
| Apomixis | 25 387 | 760 (2.91%) | 77.26% | 22.74% |
| Flowering time - total | 120 666 | 2 889 (2.34%) | 80.44% | 19.56% |
| - exons | 66 638 | 1 639 (2.39%) | 80.42% | 19.58% |
| - introns | 54 028 | 1 250 (2.26%) | 80.46% | 19.54% |
| Heat resistance - total | 81 582 | 2 263 (2.73%) | 79.22% | 20.78% |
| - exons | 51 576 | 1 399 (2.66%) | 78.77% | 21.23% |
| - introns | 30 006 | 864 (2.84%) | 79.89% | 20.11% |
| Pollen development - total | 272 591 | 7 416 (2.66%) | 79.54% | 20.46% |
| - exons | 173 714 | 4 637 (2.60%) | 79.95% | 20.05% |
| - introns | 98 877 | 2 779 (2.75%) | 78.85% | 21.15% |
| Stomatal regulation - total | 71 685 | 1 634 (2.25%) | 79.43% | 20.57% |
| - exons | 42 246 | 940 (2.19%) | 78.89% | 21.11% |
| - introns | 29 439 | 694 (2.33%) | 80.20% | 19.80% |

*: After quality filtering (mapping quality ≥20, genotype quality ≥20, read-depth ≥15). Multi-allelic (>2) polymorphic sites have been split up into separate SNVs. Percentages (in brackets) refer to the number of polymorphic sites compared to the total target length.
§: Calculated between individuals of the same ACL.

Table S5: Genetic distances between technical replicas of the same samples.

Figure S6: Principal component analysis (PCA) for different categories of loci, all samples included. Comparison of apomictic clonal lineage discrimination through principal component analysis (PCA). Different categories of loci are contrasted to reveal the difference in discriminatory power between customised (A – GBS loci; B – Custom genes) and universal (C – Angiosperms-353 + Compositae-COS) target capture sequencing. ACLs belong to Taraxacum officinale and are indicated with three-letter codes: Ala=alatum; Ekm=ekmanii; Int=interveniens; Obt=obtusifrons; Pul=pulchrifolium. Samples that can be discriminated from the core of the clonal lineage are highlighted, corresponding to sample numbering in Fig. 3 (Online Supporting Material - 'Accession Information' for more details). Scales are identical in each plot. Conserved exons from universal target capture kits provided a discrimination comparable with custom genes and GBS loci, both between and within ACLs.

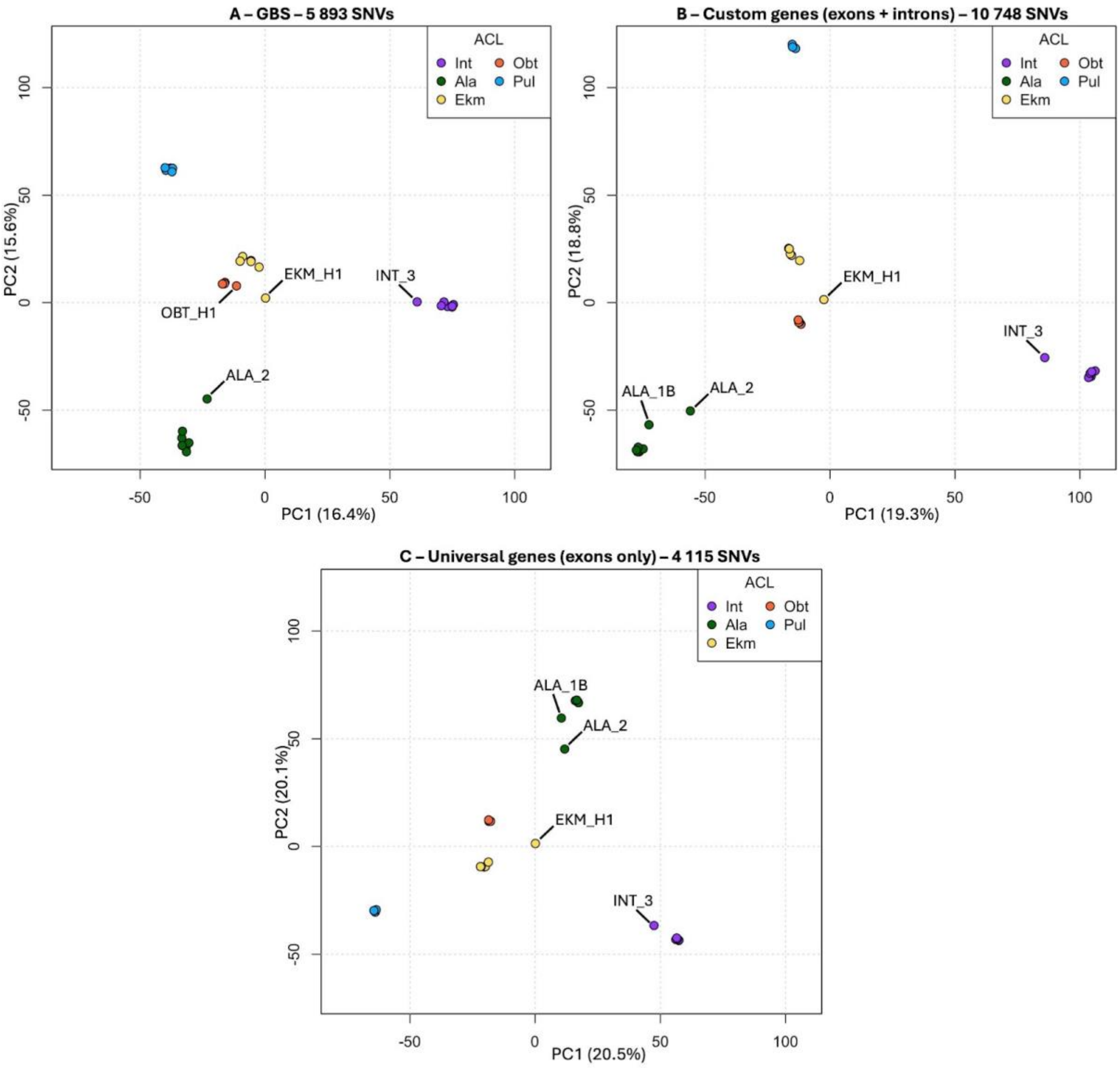
